# Metapredict enables accurate disorder prediction across the Tree of Life

**DOI:** 10.1101/2024.11.05.622168

**Authors:** Jeffrey M. Lotthammer, Jorge Hernández-García, Daniel Griffith, Dolf Weijers, Alex S. Holehouse, Ryan J. Emenecker

## Abstract

Intrinsically disordered proteins and protein regions (collectively IDRs) are critical in numerous cellular processes, yet we lack mature tools to predict their function. The first step towards elucidating IDR-mediated function necessitates IDR identification. While many disorder prediction methods exist, throughput and accuracy limit evolutionary-scale analyses. To bridge this gap, we developed metapredict V3, an updated version of our disorder predictor that enables evolutionary-scale disorder prediction. Metapredict V3 enables proteome-scale prediction with state-of-the-art accuracy in seconds and was developed with usability in mind. It is distributed as a web server, a Python software package, a command-line interface, and a Google Colab notebook. Here, we leverage the accuracy and throughput of metapredict V3 to predict disorder for over 20,000 proteomes to evaluate the prevalence of disorder across the kingdoms of life.

## INTRODUCTION

Although the structure-to-function relationship has been critical for understanding protein function, not all proteins or protein regions are well-described by a single three-dimensional structure^1^. In contrast to folded proteins, intrinsically disordered proteins and protein regions (collectively IDRs) lack a stable three-dimensional structure and are best characterized as an ensemble of interconverting conformations^2,3^. Despite this, IDRs are found in proteins with critical cellular functions, such as nuclear transport, DNA damage response, gene expression regulation, signal transduction, and stress response^4^. Although IDRs regulate many aspects of cellular function, their lack of stable structure means they are understudied relative to folded domains. As a result, our understanding of sequence-function relationships in IDRs remains limited.

The first step in investigating IDR function is to determine which regions of a protein are disordered. While experimental validation remains the gold standard for IDR identification, biophysical methods for defining IDRs are often time-consuming, low-throughput, and laborious, and require expensive equipment and considerable expertise^5–9^. Although not a perfect substitute for experimental validation, computational methods that predict disordered regions from amino acid sequences (“disorder predictors”) substantially reduce the barrier to identifying potential IDRs in a protein of interest.

Methods for predicting disorder from sequence have evolved and improved over the last two decades^10–16^. Current state-of-the-art predictors are generally highly accurate. However, while the accuracy of the top-ranked predictors is comparable, performance and accessibility are not^17,18^. Most top-performing predictors take minutes to hours per protein sequence and often require substantial computational resources to predict tens or hundreds of sequences. Beyond performance challenges for large-scale prediction, many predictors have complex installation procedures and/or web servers that are limited to processing a single sequence at a time. These factors have hampered the adoption of state-of-the-art tools with improved accuracy.

Here, we report metapredict V3, the latest version of metapredict, our high-throughput disorder predictor^19^. Metapredict V3 significantly improves accuracy and performance compared to our previous version. In terms of accuracy, metapredict V3 is comparable to the state-of-the-art predictors. In terms of performance, metapredict V3 enables prediction of entire proteomes in seconds (e.g., the yeast proteome in < 1 second, the human proteome in ∼4 seconds). This unprecedented accuracy and throughput opens the door to evolutionary-scale informatics using commodity hardware in minutes.

To ensure widespread adoption and usability, metapredict V3 is distributed in a range of implementations, which include (1) a more streamlined Python interface, (2) a modern web server (https://metapredict.net), (3) an easy-to-use command line tool, (4) a Docker container, and (5) a Colab notebook. Furthermore, despite significant changes to the underlying code, we have maintained complete backward compatibility with our previous version of metapredict. This means users who choose metapredict V3 can take full advantage of its accuracy and throughput improvements without rewriting complex bioinformatics pipelines.

The remainder of this manuscript is structured as follows: First, we describe how we developed metapredict V3 and assess its accuracy, speed, and resource requirements. Then, we leverage the high-throughput capabilities of metapredict to examine the prevalence of disorder throughout the kingdoms of life. Last, we discuss the usability of metapredict for users across a wide range of use cases and levels of expertise.

## RESULTS

### Metapredict version history

Before reporting the methods and results for metapredict V3, we first contextualize this manuscript’s work in relation to prior metapredict versions. The original metapredict (metapredict V1) was released in spring 2021^19^. In the spring of 2022, we reported an updated implementation (metapredict V2) that combined predicted pLDDT scores with consensus disorder scores^20^. Metapredict V2 represented a substantial improvement in accuracy over V1, but was comparable (if not slightly slower) than V1 (human proteome in ∼20-30 min). Metapredict V2 was reported in a preprint but was never submitted to a journal. In 2023, we implemented a parallelized batch prediction algorithm which was shared as metapredict V2-FF^21^. Metapredict V2-FF predictions are identical to V2 but enable substantially faster performance (human proteome in 2-3 minutes).

In this manuscript, we report two technical innovations. First, we report a newly trained network (metapredict V3) that improves accuracy over V1 and V2, as assessed across multiple independent evaluations. Second, we fully re-engineered the metapredict software package, implementing the original V1 and V2 versions alongside V3 and enabling full proteome predictions in just a few seconds across all three predictors. While the numerical values for V1 and V2 remain identical to those originally reported, the newly engineered implementations offer a ∼250x performance improvement over our original V1 implementation. Finally, to support the analysis of sequence databases that scale to hundreds of millions of sequences, metapredict now supports file streaming, enabling prediction across files that would not fit in memory when read directly.

### Combining disorder scores with AlphaFold2 pLDDT improves disorder prediction

We first released metapredict V1 based on “consensus” disorder prediction scores^19^. While integrating many predictor scores was powerful, prediction accuracy still had room for improvement. Shortly after our release, AlphaFold2 (AF2) was published, which popularized a confidence metric known as the <u>p</u>redicted <u>L</u>ocal <u>D</u>istance <u>D</u>ifference <u>T</u>est (pLDDT) score to convey the confidence associated with a given structure prediction^22,23^. Interestingly, it was observed that low pLDDT scores often – although not always – correlated with high disorder scores^19,23,24^. Given metapredict’s origin as a consensus disorder predictor, we wondered if we could combine pLDDT scores with metapredict scores to address the shortcomings of each method (**Fig. S1A, S1B**). We found that the best approach was to combine inverse pLDDT scores, normalized to 0-1, with metapredict disorder scores (**Fig. S2**). In this implementation, we used the higher value if either was above 0.5. If both the metapredict disorder score and the normalized inverse pLDDT score were below 0.5, we used the lower of the two (**Fig. S2**; see *Methods* for details).

Because our integrated approach relies on AF2 pLDDT scores, predicting disorder for sequences not in the AF2 database first requires AF2 structure prediction. To bypass this, we combined metapredict V1 scores and AF2 pLDDT scores obtained from AlphaFoldDB to generate a large dataset (455,666 sequences) of ‘combined disorder scores’. To focus on long disordered regions (at the expense of short loops – see below), we also applied a smoothing filter and a 25-residue sliding window to smooth boundaries. Finally, taking these integrated, smoothed scores, we used the deep learning toolkit PARROT^25^ to train a bidirectional recurrent neural network with long short-term memory (BRNN-LSTM) to predict these scores directly from amino acid sequences. This network (henceforth ‘metapredict V3’) recapitulates the ‘combined disorder scores’, thus removing the need to input AF2 pLDDT scores to generate disorder predictions (**Fig. S2**).

### Metapredict V3 is highly accurate

Many groups have devised ways to measure the accuracy of protein disorder predictors ^17,18,26–28^. To assess metapredict V3, we used the Disprot-PDB datasets from the first, second, and third iterations of the Critical Assessment of Protein Intrinsic Disorder Prediction (CAID1, CAID2, and CAID3) ^17,18,29^. The Disprot-PDB (renamed Disorder-PDB in CAID2) dataset consists of proteins with regions experimentally validated as disordered or structured^17,18^. Disordered regions come from the Disprot database^30^, while structured regions come from the PDB^31^. Metapredict V3 showed substantially improved accuracy over V1 and a modest improvement over V2. Across CAID1, CAID2, and CAID3, metapredict versions are among the most accurate disorder predictors, as assessed by the Matthews correlation coefficient, the gold-standard binary classification evaluation metric^32^ (**Fig. 1A-C, S3**). In addition to being more accurate overall, we found that metapredict V3 disorder plots were more interpretable than V1 plots and more clearly delineated IDRs from annotated folded domains (**Fig. 1D**, **Fig. S4**).

**Figure 1.**
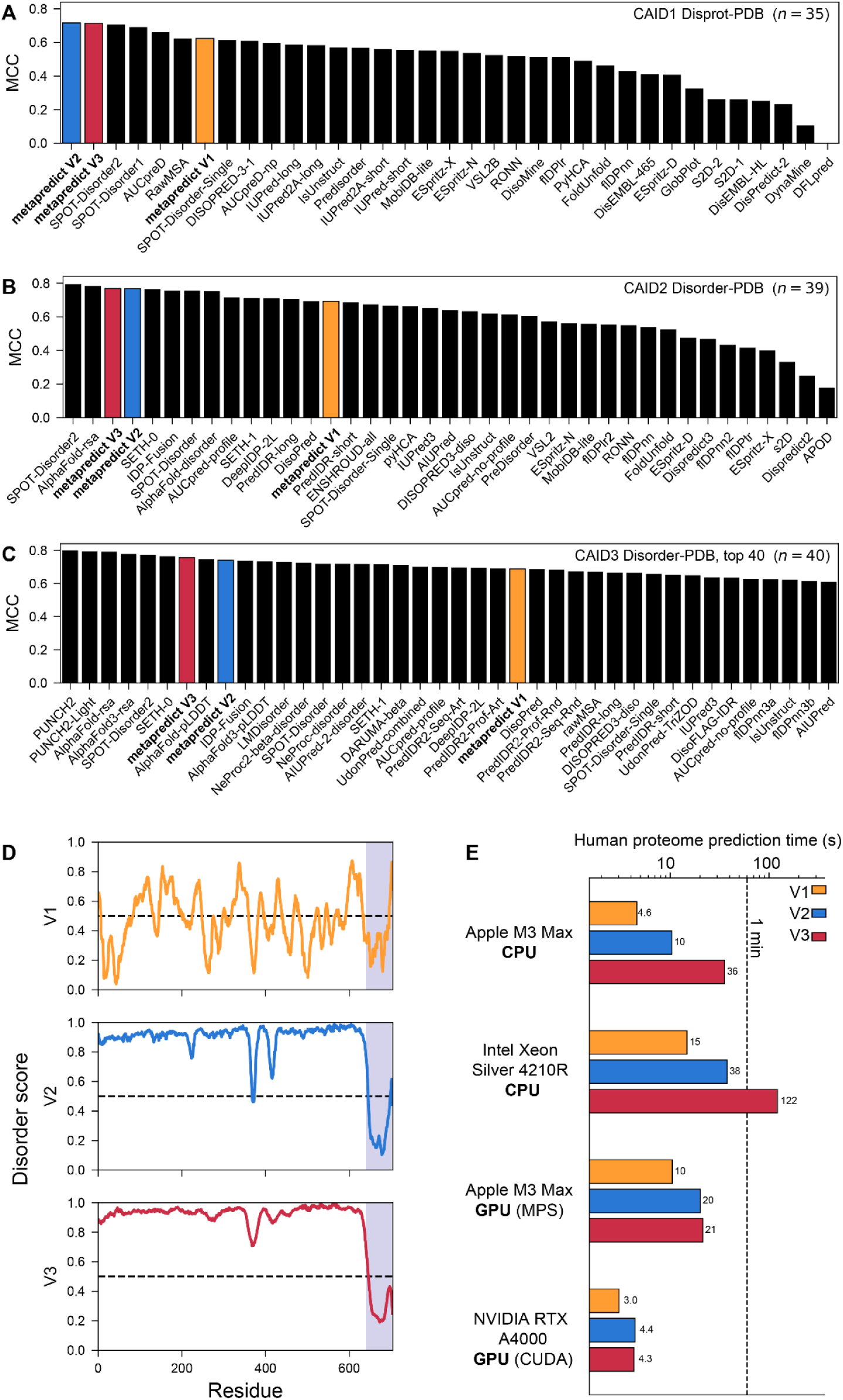
Metapredict V3 is fast and accurate. **(A–C)** Per-residue Matthews correlation coefficient (MCC) of disorder predictors on the CAID1 Disprot-PDB (A), CAID2 Disorder-PDB (B), and CAID3 Disorder-PDB (C; top 40 of 66) datasets. Metapredict V1 (orange), V2 (blue), and V3 (crimson) are highlighted. MCC needs an ordered/disordered call for every residue. For metapredict, a residue is called disordered if it lies inside a DisorderedDomain (see *Methods*). (A) Other predictors’ values are the published CAID1 MCCs. (B) We score predictors on the binary calls they submitted to CAID2. Predictors that submitted only scores are included only if a published cutoff exists (AlphaFold-rsa, AlphaFold-disorder, and RONN). (C) Predictors are scored on their submitted binary calls where available. Those that submitted only scores are cut at 0.5, except the AlphaFold2 and AlphaFold3 RSA and pLDDT tracks, which use the published AlphaFold2 cutoffs (0.581 and 0.312)<u>(Piovesan et al. 2022)</u>. Six of the predictors shown are cut at 0.5 (LMDisorder, AIUPred-2-disorder, UdonPred-combined, rawMSA, UdonPred-TriZOD and DisoFLAG-IDR), which may not be their intended cutoff. **(D)** Disorder prediction using metapredict V1 (top), V2 (middle), and V3 (bottom) for the *Saccharomyces cerevisiae* transcription factor Msn2 (UniProt ID: P33748). The region highlighted corresponds to the annotated DNA-binding domain. **(E)** Prediction speed of the reference human proteome from UniProt (one protein sequence per gene, 20,652 proteins) in seconds using metapredict V1, V2, or V3 across different hardware modalities.

### Metapredict V3 enables proteome-scale disorder prediction in seconds

In addition to a marked improvement in accuracy, we re-engineered the metapredict V3 backend to be considerably faster than our original implementation (**Fig. S5**). Specifically, we (i) enabled the prediction of disorder in batches of sequences as opposed to individual sequences, with smart padding; (ii) rewrote much of the backend code in the high-performance language Cython; and (iii) enabled improved CPU and GPU utilization.

These changes deliver substantial performance improvements compared with earlier versions of metapredict. On a single Intel CPU, the human proteome can be predicted in 15 s, 40 s, or 120 s (using metapredict V1, V2, or V3). On an Apple M3 Max chip, human proteome predictions take ∼5 s, ∼10 s, or ∼40 s (or ∼10 s, ∼20 s, and ∼20 s using MPS). Finally, on an NVIDIA A4000 GPU, across all three network versions, proteome prediction takes 3-4 seconds (**Fig. 1E**). Thus, metapredict V3 is not only accurate but also extremely fast, enabling disorder prediction at an evolutionary scale across thousands of proteomes in hours rather than weeks.

### Metapredict enables variable batch size and streaming-based predictions

In 2023, we released metapredict V2-FF, an implementation of the V2 network that batched sequences in a fixed batch size. This substantially improved performance but did not expose control of the metapredict memory footprint to users. To address this, in our V3 implementation, we rewrote batch handling, exposed batch size, and investigated optimal batch sizes across different hardware modalities to balance throughput and memory footprint (**Fig. S6-7**). This analysis allowed us to tune model- and hardware-specific defaults while exposing batch size as a variable users can modulate.

Finally, when datasets are on the order of hundreds of millions of sequences (e.g., when developing frontier protein language models), loading datasets of this size into memory can become prohibitive. To address this, we implemented a stream-based I/O interface that streams files from disk. Profiling predict_disorder_stream() against ever-increasing dataset sizes keeps memory bounded (**Fig. S8**). Here again, the chunk size – how much data is read into memory at any given moment – is exposed to users.

### Metapredict shows interpretability comparable to other top predictors

Our assessments thus far focus on accepted performance statistics commonly used to assess prediction tools, all of which confirm metapredict V3 is among the most accurate predictors (**Fig. 1, 2A, B**). However, in practice, we felt it was important to examine how well metapredict works within the actual domain of use – taking a protein sequence and identifying contiguous subregions that correctly map to disordered domains.

**Figure 2.**
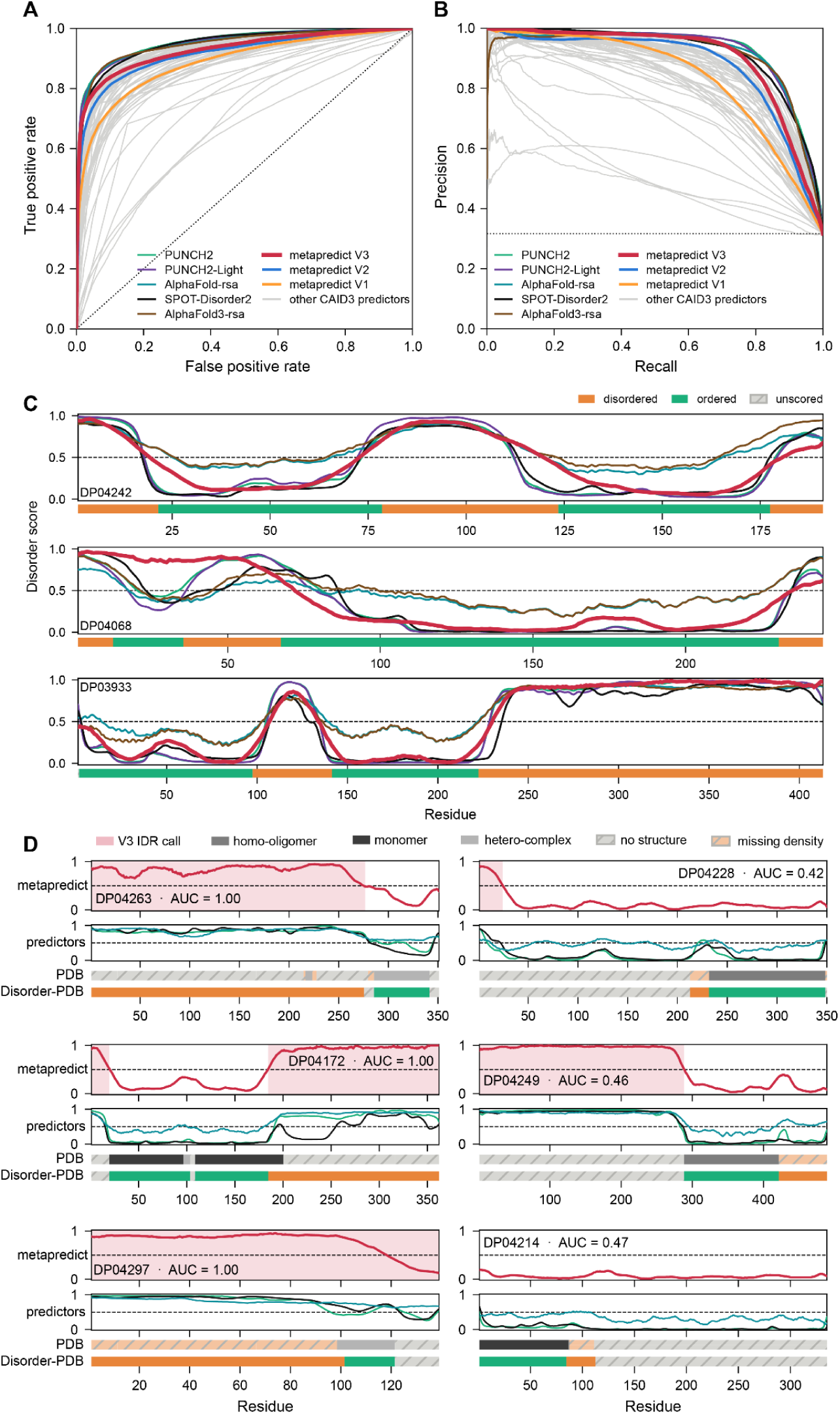
metapredict enables clear identification of contiguous disordered domains. (A) ROC and **(B)** precision–recall curves for metapredict V1, V2, and V3 and the five CAID3 disorder predictors with the highest AUC. Residues are pooled across all 319 Disorder-PDB proteins (31,401 disordered and 67,838 ordered). Grey curves are the other 58 CAID3 disorder predictors. **(C)** Per-residue disorder scores of metapredict V3 and the five top CAID3 predictors along three proteins (DisProt IDs above each trace). The bar under each trace gives the Disorder-PDB reference. **(D)** Example traces for proteins (40-500 residues) on which metapredict V3 has the highest (left) and lowest (right) per-protein AUC. Per-protein AUC is the ROC AUC of the V3 score over that protein’s scored residues, and is given above each protein after its DisProt ID.

We first compared metapredict V3 (and other top predictors) against proteins with robust coverage in Disprot-PDB, highlighting how all top predictors show a comparable ability to identify boundaries between disordered and folded domains. We noticed that in DP04068 (**Fig. 2C**, *middle*), metapredict does not identify the short 23-residue “folded” region identified at the N-terminus (residues 13-35). However, this region corresponds to a single alpha helix, which we propose is better considered a structural element within a disordered region, rather than its own ordered domain.

We also compared PDB annotations (categorizing ordered annotations as originating from monomeric proteins, homo-oligomers, hetero-complexes) with Disorder-PDB, metapredict, and several other top predictors (**Fig. 2D**). Separating out PDB data into these distinct types allowed us to investigate whether metapredict performed better or worse if putative stabilizing intermolecular interactions were excluded, although we did not find evidence for this (**Fig. S9**). With these annotations in mind, we examined proteins where metapredict performed well (**Fig. 2D**, *left*) and where it performed poorly (**Fig. 2D**, *right*). We noted two things in this visual comparison: first, places where metapredict disagreed with Disorder-PDB were often also missed by other top predictors, suggesting consistent failure modes. Second, most of the proteins where metapredict performed poorly (a small subset shown in **Fig. 2D**, right) involve short (10-49 residue) disordered regions (**Fig. S10**). Because metapredict V3 was specifically designed to identify larger contiguous disordered regions at the expense of shorter loops (driven by the sliding-window design decision), we re-assessed CAID3 accuracy by stratifying IDRS by length (**Fig. S11**). This analysis revealed that for IDRs 50-99 residues, metapredict ranked 5th by MCC, although no predictors were statistically more accurate (as assessed by bootstrap analysis), and for IDRs longer than 100 residues, metapredict ranked 4th, with only PUNCH2 as statistically more accurate.

Finally, to assess accuracy in terms of residue differences rather than rank or statistical significance, we calculated the number of correctly or incorrectly scored residues that differed between the best-scoring predictor and metapredict (**Fig. S12**). This analysis showed that the best-scoring predictor (PUNCH2) scores 1.8 more residues per 100 correctly than metapredict V3 (91.5 vs 89.6), and the ten best predictors differ by at most 2.7 residues per 100. In other words, metapredict is 1-2 residues different per 100 residues than the best-scoring predictors, and effectively statistically indistinguishable for long IDRs.

### Disordered regions are prevalent across the Tree of Life

Previous studies have found that the fraction of predicted disorder in a given proteome varies dramatically among archaea, bacteria, eukaryotes, and viruses ^33^. Given that metapredict V3 is among the most accurate available disorder predictors and can predict disorder across entire proteomes in seconds, we sought to more thoroughly characterize differences in disorder prevalence across the proteomes of these groups of organisms. To this end, we used metapredict V3 to predict disorder in 23,129 reference proteomes from the UniProt database^34^ (**Fig. 3**). Prevalence of disorder was calculated as either the average fraction of disorder per protein across a given proteome (average fraction per protein) or the total number of residues predicted to be disordered out of the total number of residues in the proteome (total fraction across proteome) for 88,082,402 protein sequences. For the results described here, we report the average fraction of disorder per protein across a given proteome; however, we note that the observed trends were similar regardless of the specific quantification method used (**Fig. S13**).

**Figure 3.**
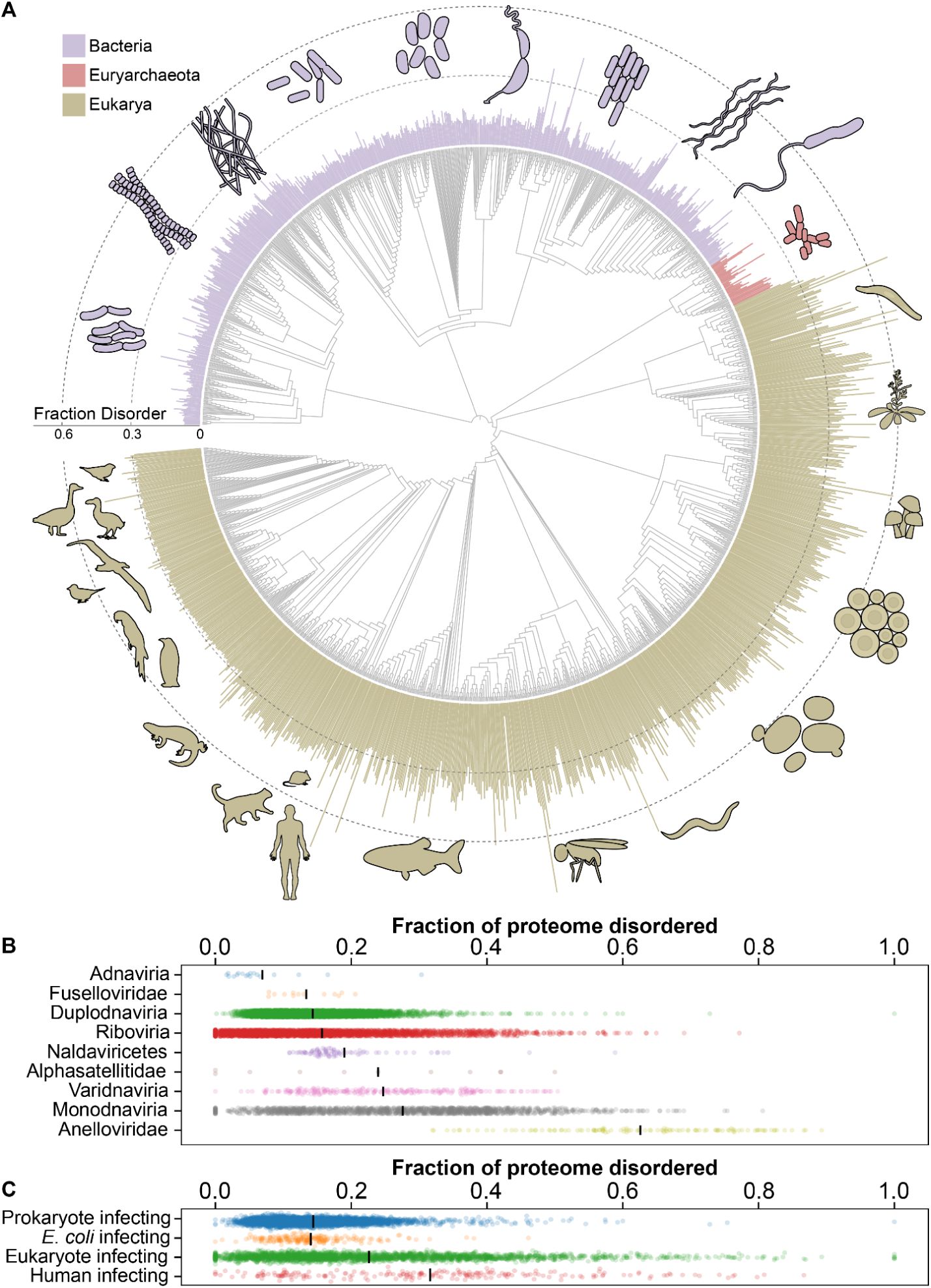
Prediction of proteomes across the kingdoms of life. **(A)** Proteome-wide disorder compared across phylogeny (12,568 proteomes) at the family taxonomic rank level (1,480 families shown). Domains are color-coded. Leaves represent families, and bars represent the average fraction of disorder across the proteome (family means). A small subset of organism images came from the NIH BIOART source (see Materials and Methods). **Fig. S13A** shows a more detailed version of this tree with major taxonomic groups. **(B)** Proteome-wide disorder fraction for viral proteomes. Virus classifications obtained from UniProt. Black lines denote the mean. **(C)** Proteome-wide disorder fraction for viruses that infect prokaryotes vs. eukaryotes. On average, viral proteomes from eukaryotic hosts are more disordered than those from prokaryotic hosts. We calculated statistical significance using the Kolmogorov-Smirnov (KS) test. Common Language Effect Size (CLES) reveals a 64% chance that a randomly selected eukaryotic virus would be more disordered than a randomly selected prokaryotic virus (see *Materials and Methods*). We also show a specific organismal comparison between *E. coli* (a prokaryote) and *H. sapiens* (a eukaryote).

To investigate the relationship between organism classification and disorder prevalence, we analyzed the evolutionary patterns of average disorder across the proteomes of Bacteria, Euryarchaeota, and Eukarya. Consistent with prior observations, we found that, on average, prokaryotes exhibited the smallest fraction of disorder among cellular organisms, in contrast to a higher fraction in most eukaryotic proteomes (**Fig. 3A**). Digging deeper, we found that among bacteria, families in the Actinobacteria (e.g., *Streptomyces*, *Mycobacterium tuberculosis*) and the PVC superphylum (Planctomycetota, Verrucomicrobiota, and Chlamydiota) tended to have a greater fraction of disorder than other bacterial groups. Similarly, when examining Euryarchaeota families, Halobacteria families frequently contained a higher fraction of disorder in their proteomes than other Euryarchaeota. The most conspicuous trend within eukaryotic lineages is the relatively low but invariable fraction of disorder among all bird proteomes (Aves), regardless of their family or taxonomic rank.

We also examined disorder prevalence across 10,561 viral proteomes. In contrast to Bacteria, Euryarchaeota, and Eukarya, the viral groups we examined (realms, classes, or families, depending on the lineage) showed wide variation in average disorder, both between groups and sometimes within them (**Fig. 3B**). Interestingly, while viral proteomes exhibit highly variable disorder fractions, there appears to be a trend of lower disorder among viruses that infect bacterial or archaeal hosts and a higher prevalence of disorder in viral proteomes whose hosts are eukaryotes (**Fig. 3C**). Further analysis of the proteomes of viruses that infect the prokaryote *E. coli* (proteome fraction disorder = 0.078) or the eukaryote *H. sapiens* (proteome fraction disorder = 0.378) again showed a correlation between viral host disorder and the fraction of disorder in a viral proteome, suggesting that the prevalence of disorder in a viral proteome converges with that of its host (**Fig. 3C**). However, further analysis of the fraction of disorder in individual viral proteomes and in the proteomes of their respective hosts is needed to establish whether this trend is biologically significant.

## DISCUSSION

Fast, accurate prediction of protein disorder is a prerequisite for systematically exploring IDRs across the Tree of Life. To address this challenge, we present metapredict V3, an accurate, performant, and broadly distributed disorder predictor.

As shown, metapredict achieves accuracy comparable to best-in-class disorder predictors. However, the top ∼10 predictors are also approximately equivalent in “real-world” accuracy (**Fig. S12)** and for IDRs of 50 residues or longer, metapredict is almost statistically indistinguishable from higher-ranked predictors (**Fig. S11**). Overall, we propose that for large-scale proteome-scale prediction tasks, the top-scoring predictors in CAID3 are approximately equally good, so throughput and scaling to large (GBs to TBs) datasets become key differentiators.

Metapredict V3 offers a substantial performance improvement over our V1 implementation. In the metapredict V1 manuscript ^19^, we reported predicting the human proteome in ∼21 minutes. With our rewrite, we can run the same proteome-scale predictions with the metapredict V1 network in 3-15 seconds (depending on hardware modality). For our most accurate network (V3), human proteome predictions take between 4 (GPU) and 120 seconds (Intel CPU). Finally, we provide metapredict implemented as a Python library, a command-line tool, various Colab notebooks, and a modern, fast web application (**Fig. S14**).

While metapredict was trained on naturally occurring sequences, we verified that metapredict V3 could accurately predict folded and disordered regions in non-natural proteins by comparing its predictions for 63 synthetic sequences (**Fig. S15**) ^35–38^. Encouragingly, metapredict V3 correctly predicts 61/63. Given the growing interest in designing and applying synthetic disordered proteins, and the limitations of language-model-based approaches when moving out of distribution ^39,40^, verifying disorder predictors work on “out-of-distribution” sequences will likely become increasingly important.

Metapredict V3’s accuracy, performance, and ease of use open the door to large- or small-scale bioinformatics with limited resources. For example, we systematically examined the 1608 transcription factors in the human proteome to investigate disorder and composition across transcription factor IDRs (**Fig. S16**)^41^. This analysis revealed that TFs are highly disordered, with TF median disorder at 57% (compared to 24% for the median human protein). Critically, what was previously an involved, time-consuming exercise can now be done in minutes with a few lines of code. Finally, our IDRome Google Colab notebook enables systematic prediction and annotation of all IDRs in a FASTA file within minutes, opening the door to proteome-scale IDR exploration with a web browser ^21^.

In conclusion, metapredict V3 combines state-of-the-art accuracy in disorder prediction with best-in-class performance and broad distribution across multiple modalities. Its performance enables evolutionary-scale disorder prediction, revealing the breadth of disorder propensities across the Tree of Life. We anticipate metapredict V3 facilitating the investigation of intrinsically disordered regions across many systems.

## ACKNOWLEDGEMENTS

Metapredict V3 would not be possible without the extraordinary amount of work carried out by developers of the various dependencies that metapredict requires to function, including Cython, PyTorch, NumPy, Matplotlib, Scipy, urllib3, tqdm, and PyTorch-Lightning. We also thank the entire Tosatto group, the ELIXIR Intrinsically Disordered Proteins Community, and HUPO-PSI Intrinsically Disordered Proteins Community (notably Silvio Tosatto, Zsuzsanna Dosztanyi, Damiano Piovesan, Wim Vranken, and Norman Davey) for all the European-funded bioinformatics work that largely fuels the international intrinsically disordered proteins informatics space. We also thank Danilo dos Santos Pereira for help with the phylogenetic analyses. We thank Patrick Kunzmann, joushvak17, members of the Freid lab, and Henrik Åhl for code contributions to metapredict. We also thank Emery Usher and Jackie Pelham for their helpful comments on this manuscript.

This work was supported by a Research Grant from the Human Frontiers Research Program (HFSP) grant RGP0015/2022 to A.S.H and D.W. JML was supported by the National Science Foundation via grant number DGE-2139839 and by the Frontera Computational Sciences Fellowship. D.G. was supported by the NSF via award no. DGE-2139839 and the Millipore Sigma Fellowship.

## AUTHOR CONTRIBUTIONS

**CRediT statement: JML** (Conceptualization, Methodology, Software, Formal analysis, Validation, Investigation, Data Curation, Writing - Original Draft, Funding acquisition, Visualization), **JHG** (Methodology, Software, Investigation, Writing - Review & Editing), **DG** (Conceptualization, Software, Funding acquisition), **DW** (Funding acquisition, Project administration, Supervision), **ASH** (Conceptualization, Formal analysis, Resources, Review & Editing, Supervision, Project administration, Funding acquisition), **RJE** (Conceptualization, Methodology, Software, Formal analysis, Validation, Investigation, Data Curation, Writing - Original Draft, Funding acquisition, Visualization)

## SUPPLEMENTARY INFORMATION

**Document S1**. Figures S1-S16 as a .pdf file.

**Figure and analysis reproduction:** All code and data to reproduce figures is available at https://github.com/holehouse-lab/supportingdata/tree/master/2026/metapredict_2026

## MATERIALS AND METHODS

### Combining predicted pLDDT (ppLDDT) scores (V2) or AF2 pLDDT scores (V3) with disorder scores to make ‘combined disorder scores’

While metapredict V1 predicted disorder scores from a set of existing disorder predictors, metapredict V2 and V3 integrate these predictions with either predicted (V2) or actual (V3) pLDDT (predicted local distance difference test) obtained from AlphaFold2. pLDDT scores represent confidence in very local structural predictions from a structure prediction model (in our case, AlphaFold2).

The V2 and V3 networks predict the composite disorder/(p)pLDDT score. The disorder scores were combined with the (p)PLDDT (predicted in V2, actual in V3) as follows:

1. pLDDT (V3) or ppLDDT (V2) scores were divided by 100 and then were normalized using a base value equal to 0.35 and a top value equal to 0.95 such that final values range between 0 and 1.0
2. pLDDT or ppLDDT scores were inverted such that a higher value corresponded to possible disordered regions and a lower value corresponded to possible folded regions,
3. For each residue in a given protein, we compared the scaled and normalized ppLDDT (V2) or AF2 pLDDT (V3) scores with metapredict V1 disorder scores. If either value was above 0.5, we used the higher value. If both values were below 0.5, we used the lower value.
4. We applied a Savitzky-Golay filter to the resulting scores to smooth them.
5. Any scores above 1 were clipped to 1.
6. Any scores below 0 were clipped and set to 0
7. For metapredict V3 combined scores only, we applied an additional smoothing step using a moving average across a 25-residue window to the final scores.

To generate the metapredict V2 scores, ‘version 1’ of the pLDDT predictor was used to create ppLDDT scores. The code for generating the hybrid scores can be found in the GitHub repo at metapredict/backend/creating_V2_and_V3_scores.py with the *meta_predict_hybrid()* function for V2 scores and the *meta_predict_hybrid_v3()* function for V3 scores.

### Sequences used to train the metapredict V2 network

363,265 sequences from 21 proteomes were used to train metapredict V2, as listed below: UP000002485_284812_SCHPO,UP000000805_243232_METJA, UP000001450_36329_PLAF7, UP000005640_9606_HUMAN, UP000001584_83332_MYCTU, UP000001940_6239_CAEEL, UP000000625_83333_ECOLI, UP000002296_353153_TRYCC, UP000000803_7227_DROME, UP000007305_4577_MAIZE, UP000002195_44689_DICDI, UP000002311_559292_YEAST, UP000002494_10116_RAT, UP000008153_5671_LEIIN, UP000008816_93061_STAA8, UP000006548_3702_ARATH, UP000008827_3847_SOYBN, UP000000437_7955_DANRE, UP000000589_10090_MOUSE, UP000059680_39947_ORYSJ, UP000000559_237561_CANAL

### Sequences used to train the metapredict V3 network

The sequences used to train the metapredict V3 network came from the Swiss-Prot database ^42^, which was filtered to remove redundant sequences and then further filtered for sequences with corresponding AlphaFold2 ^23^ structures, because AF2 pLDDT scores are required to generate the V3 ‘combined disorder scores.’ This resulted in a final set of 455,666 sequences that were ultimately used for training.

### Training metapredict V2 using PARROT

Using PARROT^25^, we trained a bidirectional recurrent neural network with long short-term memory (BRNN-LSTM) on the metapredict ‘combined disorder scores’ created by combining ppLDDT and metapredict V1 disorder scores. We trained this network as a regression model using one-hot encoding, a hidden vector size of 10, and 2 layers. We also used a learning rate of 0.001, a batch size of 32, and trained for 200 epochs. We split the 363,265 sequences from the 21 proteomes 70:15:15 for training, validation, and testing, respectively.

### Training metapredict V3 using PARROT-lightning

We leveraged an adapted version of PARROT, parrot-lightning (https://github.com/idptools/parrot/tree/parrot-lightning), to train a similar BRNN-LSTM architecture on metapredict hybrid scores created from a deduplicated set of protein sequences from the SWISSPROT sequence database. The resulting dataset contained 455,666 unique sequences. As in our previous training schemes, we split our data into 70:15:15 for training, validation, and testing, respectively.

To train metapredict V3, we used this data and the optimize_params.py script in parrot-lightning to perform Bayesian hyperparameter optimization with Optuna^43^. Specifically, we searched for optimal learning rates and hidden sizes that conferred the lowest validation loss on our validation set. Our final network had 2 BRNN-LSTM layers, with hidden sizes of 52. For model training, we used stochastic gradient descent with Nesterov momentum (momentum=0.9968), a learning rate of 0.01427, and a batch size of 256. Training used automatic mixed precision and a gradient clipping value of 1.

### Usage and features

Metapredict can be used in five different ways.

#### Python API and command-line tool

You can download and run metapredict locally, where you can use it from the command line or as a Python module. The command-line interface includes functionality to predict disorder directly from sequence (per-residue scores or contiguous disordered regions), either from a passed-in sequence or an input FASTA file. The command-line tool also fetches sequences directly from UniProt using the UniProt accession number or the common organism and protein name. Finally, in addition to predicting disorder, the command-line tool also enables per-residue disorder plotting.

The Python module includes all the functionality described above, can also predict per-residue pLDDT scores, and offers substantial customizability in various parameters that may be of interest (e.g., thresholds for calling something disordered, minimum IDR size, minimum folded domain size, etc.). The command-line interface and Python module automatically detect CUDA-enabled GPUs and will run predictions on the GPU if available. More detailed documentation can be found at https://metapredict.readthedocs.io/en/latest/.

#### Webserver

Metapredict can be used by a web server to run predictions and visualize IDRs at https://metapredict.net/. The default version for the metapredict server is currently V3, although alternative versions can be selected using the toggle switch.

#### Docker container

Metapredict is also distributed as a Docker container (https://hub.docker.com/r/holehouselab/metapredict), enabling large-scale predictions without having to install or manage dependencies.

#### Colab notebook

Finally, metapredict can be used in a Google Colab notebook to enable proteome-wide predictions from FASTA files without having to install metapredict locally (https://colab.research.google.com/github/idptools/metapredict/blob/master/colab/metapredict_colab.ipynb). Metapredict V3 is also included as the default predictor for the IDRome constructor notebook (https://colab.research.google.com/github/holehouse-lab/ALBATROSS-colab/blob/main/idrome_constructor/idrome_constructor.ipynb) which lets you predict biophysical and sequence properties for all IDRs in a proteome.

### Hardware for performance tests

We ran performance tests (**Fig. 1E**, **S5-8**) on Apple, NVIDIA, and Intel hardware. The first machine used was a 14-inch MacBook Pro (2023) with an Apple M3 Max chip (16-core CPU with 12 performance and 4 efficiency cores; 40-core GPU), 128 GB of unified memory and a 4 TB SSD, running macOS 26.5.1. GPU tests on this machine used PyTorch’s Metal Performance Shaders (MPS) backend. The second was a Linux workstation with an Intel Xeon Silver 4210R CPU (10 cores, 20 threads, 2.40 GHz), 64 GB of RAM, and an NVIDIA RTX A4000 GPU (16 GB), running Ubuntu 20.04 (Linux kernel 5.15) with NVIDIA driver 580.126.09. GPU tests on this machine used CUDA. We used Python 3.12.13 and PyTorch 2.10.0 on the MacBook Pro, and Python 3.14.8 and PyTorch 2.14.1 (CUDA 13.0 build) on the workstation.

### Performance tests using human proteome prediction

**Fig. 1E**, and **Fig. S5-7** used the human reference proteome from UniProt (UP000005640), downloaded on July 5th 2026. This dataset contains 20,652 sequences (11,453,425 residues; the longest is titin, at 34,350 residues), while **Fig. S8** used synthetic sequences (see "Synthetic sequences for memory profiling"). Unless stated otherwise, each network used metapredict’s default batch size for the device (256 sequences on the CPU and on CUDA GPUs, 512 on Apple MPS). Within a call, sequences are batched longest-first, and each sequence is padded to the length of the longest one in its batch. Before timing, an untimed warm-up prediction on a few sequences loaded the network weights and initialized the device. Times are wall-clock times (Python time.perf_counter) of the prediction call alone, excluding reading the FASTA file and the warm-up, and for GPU runs the device was synchronized before the clock was stopped so that all queued GPU work was included.

Throughput is the number of residues predicted divided by this time. Peak memory is the peak resident set size (RSS) of the Python process during the timed prediction, sampled by a background thread every 0.1 s (every 4 ms in **Fig. S8**). Peak GPU memory is torch.cuda.max_memory_allocated on CUDA GPUs, which counts memory held by tensors and excludes PyTorch’s cache, and the largest sampled value of torch.mps.driver_allocated_memory on MPS, which includes the cache. On Apple Silicon, the GPU memory allocated is not counted in the process RSS. For **Fig. 1E** and **Fig. S5**, we timed each network (V1, V2, and V3) predicting the whole proteome in a single call on each device (CPU and MPS on the MacBook Pro; CPU and CUDA on the workstation), once per configuration, each in a separate Python process.

For **Fig. S6**, each network predicted the whole proteome on each device at batch sizes of 32, 64, 128, 256, 512 and 1,024, with cached GPU memory released before each measurement. Batch sizes that ran out of memory (V3 at 512 and 1,024 on the RTX A4000) have no time. Unrounded scores at each batch size differed from those at the smallest batch size by at most 6.1 × 10⁻⁶, confirming that batch size does not change predictions. We ran CPU sweeps three times on each computer (median plotted) and GPU sweeps once. For **Fig. S7**, we drew random subsets of 10, 50, 100, 200, 500, 1,000, 2,000, 3,000, 4,000, 5,000, 8,000, 10,000, and 15,000 sequences from the proteome. We timed each subset as a single batched call on the list of sequences and as a loop of single-sequence calls. Because metapredict always predicts a single sequence on the CPU, sequential times are CPU times. Batched calls used each network’s default device on the MacBook Pro (CPU for V1 and V2, MPS for V3) and were run once on the CPU and once on the GPU on the workstation, whose sequential times are the median of these two runs. One random generator was shared across the three networks, so V1, V2 and V3 were timed on different subsets, but each network was timed on the same subsets on both computers. The hardware, operating system, Python, PyTorch, and metapredict version (with git commit) for every measurement are recorded in the benchmark results files.

### Performance tests using randomly generated sequences

The synthetic sequences used for memory profiling (**Fig. S8**) were generated by pulling sequence lengths from a log-normal distribution (μ = 5.7, σ = 0.55 on the natural-log scale; median ∼300 residues), rounded down to whole residues, and limited them to 40–2,500 residues. We drew each residue at random from the 20 standard amino acids. We generated seven datasets of 20,000, 80,000, 250,000, 500,000, 720,000, 1,440,000 and 2,880,000 sequences, containing 7.0 million to 999 million residues (mean length 347 residues; the largest dataset is about 87 times the size of the human proteome). We re-seed the generator for each dataset, so each dataset is identical across machines. We used this approach instead of our IDR design package GOOSE to ensure the sequences tested here were truly random for performance purposes, rather than biasing them toward disordered tendencies. We predicted each dataset with metapredict V3 using predict_disorder and predict_disorder_stream, both at a smaller-than-default batch size of 32, with a chunk size of 20,000 for streaming. Each measurement ran in a fresh Python process and recorded peak resident memory and total run time. There was one measurement per dataset, function, and device.

### CAID1, CAID2, and CAID3 analyses of metapredict

We assessed the three metapredict networks on the Disprot-PDB dataset from CAID1 and the Disorder-PDB datasets from CAID2 and CAID3. For each round, we predicted the CAID target sequences with metapredict (V1, V2, and V3 networks) using predict_disorder_caid(), rather than using the metapredict predictions distributed by CAID. To quantify accuracy, we considered a residue predicted disordered if it fell within a DisorderedDomain, defined using metapredict’s default IDR-calling algorithm (minimum_IDR_size = 12, minimum_folded_domain = 50, gap_closure = 10). We calculated MCC over all residues of all targets with a reference label; we excluded unlabelled residues. For CAID1 (**Fig. 1A**), we used the MCC values for the other predictors published by CAID1. For CAID2 and CAID3 (**Fig. 1B, 1C** and **S12**), we recomputed every predictor’s MCC from the per-residue prediction files distributed by CAID, scoring each predictor on its submitted binary calls where available. For CAID2, we included predictors that submitted only scores only if a published disorder cutoff exists (AlphaFold-rsa 0.581, AlphaFold-disorder 0.312, and RONN 0.5), yielding 39 predictors, including metapredict. For CAID3, we included all 66 disorder predictors. We assigned binary calls to the 19 predictors with only submitted scores at a cutoff of 0.5, except for the AlphaFold2 and AlphaFold3 RSA and pLDDT tracks, which used AlphaFold2 cutoffs of 0.581 and 0.312. Fifteen predictors were cut at 0.5: LMDisorder, AIUPred-2-disorder, UdonPred-combined, UdonPred-TriZOD, UdonPred-DisProt, rawMSA, DisoFLAG-IDR, RONN, ESMDisPred-1, ESMDisPred-2, ESMDisPred-2PDB, DisEMBL-dis465, DisEMBL-disHL, DisoMine and OPAL. A cutoff of 0.5 is not necessarily what these methods were designed for, and it can substantially lower their MCC. For example, ESMDisPred-2PDB has an MCC of 0.756 at its Fmax threshold (**Fig. S3D**) but 0.506 at 0.5. The threshold-free metrics in **Fig. 2** and S3 (AUC, APS and Fmax) do not depend on these cutoffs.

### CAID3 accuracy metrics

For **Fig. 2A,B** and **Fig. S3, S9** and **S11**, we scored the 63 CAID3 disorder predictors (binding and linker predictors excluded) and the three metapredict networks, 66 in total, on the CAID3 Disorder-PDB dataset (319 proteins). We used the per-residue scores each predictor submitted to CAID3 and metapredict. Residues annotated as disordered in DisProt count as disordered (31,401), residues resolved in the PDB count as ordered (67,838), and all other residues are excluded. As in the CAID assessments, we pool residues across proteins, and each predictor is scored only on the proteins it predicted (for example, SPOT-Disorder2 covers 298 of the 319). We calculated the area under the ROC curve (AUC), the average precision score (APS), and Fmax (the highest F1 score over all score cutoffs) with scikit-learn, and the MCC at the cutoff that gives Fmax.

### Per-protein AUC and PDB structural evidence

Per-protein AUC (Fig. 2D) is the ROC AUC of the metapredict V3 score over one protein’s scored residues. We considered proteins of 40–500 residues with at least 15 disordered and 15 ordered scored residues (116 proteins). Of these, 27 have an AUC of 1.00, and we ranked them by their number of disordered residues. Fig. 2D shows three of the five best and three of the five worst proteins. To show the structural evidence behind each residue, we mapped each protein from DisProt to UniProt, found every PDB entry covering it with PDBe SIFTS, and took the residues modeled in each entry from PDBe polymer coverage (accessed Oct. 2026). We classed each entry as a monomer (one copy of the protein and no other polymer in the deposited model), a homo-oligomer (several copies and no other polymer), or a hetero-complex (anything else, including entries whose composition could not be retrieved). Each residue was labeled by the first of these that applies: modeled in a monomer, modeled in a homo-oligomer, modeled only in a hetero-complex, inside a construct but not modeled ("missing density"), or not in any PDB construct. Residues modeled in NMR and cryo-EM structures count as modeled. For the PDB-disorder-monomer benchmark (**Fig. S9**), residues modeled in at least one monomer entry are ordered, residues inside a monomer construct but modeled in none are disordered, and all other residues are excluded (134 proteins; 32,904 ordered and 3,807 disordered residues). These labels come from the PDB alone, so 835 of the 3,807 "disordered" residues are modeled in a homo-oligomer or complex.

### Scoring by IDR length

For **Fig. S11**, an IDR is a maximal run of residues labeled disordered in CAID3 Disorder-PDB. Unscored residues end a run, so some short IDRs are the scored ends of longer disordered regions; the shortest IDR is 10 residues. We put IDRs into four bins: 10–19, 20–49, 50–99, and ≥100 residues (158, 135, 68, and 94 IDRs). Each bin retains all 67,838 ordered residues and only the disordered residues in that bin. We scored each bin as in **Fig. S3**, choosing the Fmax cutoff separately for each bin. To test whether a predictor ranked above metapredict V3 is significantly more accurate (**Fig. S11E,F**), we used a paired bootstrap over proteins. We drew 1,000 resamples of the 319 proteins with replacement and used the same resamples for every predictor. In each resample, we scored the predictor and V3 on the drawn proteins the predictor covers. We mark a predictor if the 2.5th percentile of its difference from V3 is above zero.

### Short-IDR examples

For **Fig. S10**, short IDRs are IDRs (defined as for **Fig. S11**) of 10–49 residues. Metapredict V3 misses a short IDR when fewer than half of its residues score at least 0.5 and fewer than half lie in a predicted IDR. We classed each missed IDR as a terminal tail if it reaches either terminus, as a loop if it has ordered residues on both sides, and as next to unscored residues otherwise. We selected the ten proteins with the most missed short-IDR residues, each with at most 600 residues. Each class was represented in proportion to its share of all missed short-IDR residues (five loops, three next to unscored residues, two tails), and we classify a protein by the context of most of its missed residues. The pLDDT track is CAID3’s AlphaFold-pLDDT prediction, which CAID distributes as 1 − pLDDT/100^44^; we converted it back to pLDDT. Structures are AlphaFold DB models (AF-[UniProt]-F1, version 6, downloaded October 2026)^45^, whose sequences match the CAID3 sequences. We rendered structures in PyMOL 3.2 (open-source) and colored them by standard pLDDT.

### Average fraction of disorder across proteome comparisons

To compare the average fraction of disorder across viruses, bacteria, eukaryotes, and archaea, we used 23,129 of the 23,159 available UniProt reference proteomes. For this analysis, we used the UniProt API to download the proteomes; 30 of the 23,159 proteomes failed to download and were therefore excluded from the final analysis. The analysis included 10,561 proteomes classified as ‘viruses’, 9,592 as ‘bacteria’, 2,616 as ‘eukaryota’, and 360 as ‘archaea’. For **Fig. 3B** and **Fig. S13B**, we further required each viral proteome to contain at least two proteins (10,014 viral proteomes) and each viral group to contain at least 10 proteomes. To be included in **Fig. 3B** or **Fig. S13B**, a viral group had to have at least 10 proteomes. For **Fig. 3C** or **Fig. S13C**, we included only viruses with identified hosts in Virus-Host DB^46^.

We identified disordered regions using the metapredict DisorderedObject, obtained with the metapredict function predict_disorder() with return_domains=True. For each protein, we identified disordered domains using the object variable .disordered_domains. To quantify average disorder per protein in each proteome, we divided each protein’s total IDR length by its total length. Then, we averaged these fractions to obtain the average disorder per protein across the entire proteome. Because this approach does not account for different protein lengths, in that a 10,000-amino-acid-length hypothetical protein that is fully disordered counts towards the total average equally as a 40-amino-acid-length protein, we also calculated the ‘total disorder’ in each proteome. We calculated ‘total disorder’ by summing the number of amino acids annotated as IDRs across the entire proteome and dividing by the total number of amino acids in the proteome. For **Fig. 3**, we use the average disorder per protein across each proteome. We note no major differences between the two methods, and we include a version of **Fig. 3** using ‘total disorder’ as **Fig. S13**.

A subset of organisms shown for illustration in **Fig. 3A** were from the NIH BIOART repository. Specifically, the following images were used:

1. Ryan Kissinger. (2024) Bacillus Bacteria. NIH BIOART Source. https://bioart.niaid.nih.gov/bioart/43
2. Ryan Kissinger. (2024) Spirochete. NIH BIOART Source. https://bioart.niaid.nih.gov/bioart/496
3. Ryan Kissinger. (2024) Microbiota. NIH BIOART Source. https://bioart.niaid.nih.gov/bioart/349
4. Ryan Kissinger. (2024) Plasmodium. NIH BIOART Source. https://bioart.niaid.nih.gov/bioart/414
5. Ryan Kissinger. (2024) Lizard Outline. NIH BIOART Source. https://bioart.niaid.nih.gov/bioart/302

### Statistical testing

To determine whether the differences observed in the average fraction of disorder in the proteomes of viruses that infect prokaryotes versus viruses that infect eukaryotes or in the average fraction of disorder in the proteomes of viruses that infect *E. coli* versus viruses that infect *H. sapiens* (**Fig. 3C**) were statistically significant, we first ran a Shapiro-Wilk test to determine whether we could use a statistical test that assumes a normal distribution of data. We found that none of the datasets were normally distributed (all P-values < 1e^-4^). Therefore, we used a 2-sample Kolmogorov-Smirnov test, which does not assume normality. Both comparisons were statistically significant (P-value < 1e^-16^), with the comparison between viruses that infect prokaryotes and those that infect eukaryotes having a D statistic of 0.311, and the comparison between viruses that infect *E. coli* and those that infect *H. sapiens* having a D statistic of 0.581. Next, we used a Mann-Whitney U test to determine whether the median disorder differed between groups in our two comparisons. We found that the differences in median fraction of disorder between viruses that infect prokaryotes versus those that infect eukaryotes, and between viruses that infect *E. coli* versus those that infect *H. sapiens,* were statistically significant (P-value < 1e^-16^). Finally, we calculated the Common Language Effect Size (CLES) for our two comparisons. Comparing the fraction of proteome disorder for eukaryote-infecting viruses versus prokaryote-infecting viruses yielded a CLES of 0.6415, suggesting a 64.15% chance that a randomly selected virus from the eukaryote-infecting group would have a higher fraction of proteome disorder than a randomly selected virus from the prokaryote-infecting group. In comparing *H. sapiens*-infecting viruses to *E. coli*-infecting viruses, the CLES value was 0.7742, suggesting a 77.42% chance that a randomly selected virus from the H. sapiens-infecting group would have a higher fraction of proteome disorder than a randomly selected virus from the *E. coli*-infecting group. We carried out all statistical tests using SciPy ^47^.

### Comparison with synthetic proteins

Synthetic proteins were taken from four papers and analyzed using metapredict to predict folded and disordered regions^35–38^. The fraction disordered was calculated based on the fraction of the residues that fell within a metapredict-defined disordered domain.

### Transcription factor analysis

The 1608 human transcription factors list was taken from Lambert *et al.* ^41^ and analyzed using metapredict to predict folded and disordered regions. The fraction disordered was calculated as the fraction of residues that fell within a metapredict-defined disordered domain. We obtained human proteome and TF sequences on Oct 29th, 2024, from UniProt^34,42^.

### Phylogenetic analysis

The taxize package^48^ was used to obtain taxon information from the species in the proteomes list and to draw a consensus universal phylogenetic tree. After retrieving data with taxize, we extracted family/species information and calculated, per family, either the mean predicted fraction of disorder in the total proteome or the average disorder per protein across the proteome. See the “Average fraction of disorder across proteome comparisons” section for details on calculating the average fraction of disorder. We drew the universal tree with Taxize using a single-species NCBI ID per family to build the species tree. We homogenized tree branch sizes and plotted family-predicted mean data onto the tree using iTOL^49^. We manually edited the final tree drawings to produce the final figures. R code for the analysis can be found in the associated code repository.

### Web server implementation

metapredict.net is a Python web application built with Flask (v3) and served in production by Gunicorn behind an nginx reverse proxy on a CPU-only Linux server. Predictions use metapredict running on a CPU-only build of PyTorch. Users can choose any of the three metapredict disorder networks (V1, V2 and V3; V3 is the default) and either of the two predictors of AlphaFold2 pLDDT scores (V1 and V2; V2 is the default). Sequences can be pasted as plain text or FASTA, or fetched by UniProt accession using the UniProt REST API. The server cleans each input before prediction. It removes FASTA headers and whitespace, converts non-standard residues to a standard equivalent (B→N, U→C, X→G, Z→Q, J→L, O→K), removes any other characters (and tells the user which ones), and checks the result with protfasta. Inputs are limited to 30,000 residues.

Metapredict’s domain-decomposition routine defines disordered and folded domains. Users can change the disorder threshold (default 0.5 for V2 and V3, 0.42 for V1), minimum IDR length (12 residues), minimum folded domain length (50 residues) and gap closure (10 residues). For each predicted IDR, the server reports mean disorder, the fraction of charged residues (FCR), net charge per residue (NCPR)^50^, charge patterning (κ)^51^, hydropathy^52^, and the fractions of polar, aromatic, and aliphatic residues^53^. FCR, NCPR, κ, and hydropathy are computed with SPARROW (https://github.com/idptools/sparrow). Users can also select any region of the sequence interactively. Selected regions additionally report molecular weight, proline fraction, and isoelectric point. The isoelectric point is found by bisecting on pH, applying the Henderson-Hasselbalch equation with EMBOSS side-chain pKa values. For regions of at least 15 residues, ALBATROSS^21^ predicts ensemble dimensions: radius of gyration (R_g_) and end-to-end distance (R_e_) for IDRs, plus the scaling exponent and asphericity for selected regions. R_g_ and R_e_ are also reported divided by the expectation from the analytical Flory random coil (AFRC) model^54^, so a value of 1 means the dimensions match an ideal chain.

The front end is a single-page interface written in plain JavaScript, HTML, and CSS with no third-party libraries. The server returns each prediction as JSON, and the browser builds all downloads from that response: per-residue scores (CSV), IDR sequences (FASTA), and figures (PNG and SVG). Each sequence is therefore predicted only once. Disorder and pLDDT profiles come from a single set of drawing instructions rendered either to an HTML canvas (for PNG) or to SVG, so the two figure formats are identical. A simplified view (/simple) shows only the disorder profile and IDRs, using default settings. To keep response times consistent, each Gunicorn worker runs every disorder, pLDDT and ALBATROSS network once on a test sequence before it accepts requests, so no user request has to wait for a model to load. PyTorch’s oneDNN backend is turned off for LSTM inference. These networks are small, so there is no speed cost, and this avoids intermittent oneDNN errors when a worker runs a network for the first time.

### Code availability

The code for metapredict is available at https://github.com/idptools/metapredict. The code used for the generation of figures and for benchmarking accuracy/performance can be found at https://github.com/holehouse-lab/supportingdata/tree/master/2026/metapredict_2026.

## REFERENCES

1. Wright, P. E. & Dyson, H. J. Intrinsically unstructured proteins: re-assessing the protein structure-function paradigm. J. Mol. Biol. 293, 321–331 (1999).

2. van der Lee, R., Buljan, M., Lang, B., Weatheritt, R. J., Daughdrill, G. W., Dunker, A. K., Fuxreiter, M., Gough, J., Gsponer, J., Jones, D. T., Kim, P. M., Kriwacki, R. W., Oldfield, C. J., Pappu, R. V., Tompa, P., Uversky, V. N., Wright, P. E. & Babu, M. M. Classification of intrinsically disordered regions and proteins. Chem. Rev. 114, 6589–6631 (2014).

3. Tompa, P. Intrinsically unstructured proteins. Trends Biochem. Sci. 27, 527–533 (2002).

4. Holehouse, A. S. & Kragelund, B. B. The molecular basis for cellular function of intrinsically disordered protein regions. Nat. Rev. Mol. Cell Biol. 25, 187–211 (2024).

5. Cubuk, J., Stuchell-Brereton, M. D. & Soranno, A. The biophysics of disordered proteins from the point of view of single-molecule fluorescence spectroscopy. Essays Biochem. 66, 875–890 (2022).

6. Camacho-Zarco, A. R., Schnapka, V., Guseva, S., Abyzov, A., Adamski, W., Milles, S., Jensen, M. R., Zidek, L., Salvi, N. & Blackledge, M. NMR provides unique insight into the functional dynamics and interactions of intrinsically disordered proteins. Chem. Rev. 122, 9331–9356 (2022).

7. Kikhney, A. G. & Svergun, D. I. A practical guide to small angle X-ray scattering (SAXS) of flexible and intrinsically disordered proteins. FEBS Lett. 589, 2570–2577 (2015).

8. Schuler, B., Soranno, A., Hofmann, H. & Nettels, D. Single-molecule FRET spectroscopy and the polymer physics of unfolded and intrinsically disordered proteins. Annu. Rev. Biophys. 45, 207–231 (2016).

9. Gibbs, E. B., Cook, E. C. & Showalter, S. A. Application of NMR to studies of intrinsically disordered proteins. Arch. Biochem. Biophys. 628, 57–70 (2017).

10. Walsh, I., Martin, A. J. M., Di Domenico, T. & Tosatto, S. C. E. ESpritz: accurate and fast prediction of protein disorder. Bioinformatics 28, 503–509 (2012).

11. Mészáros, B., Erdos, G. & Dosztányi, Z. IUPred2A: context-dependent prediction of protein disorder as a function of redox state and protein binding. Nucleic Acids Res. 46, W329–W337 (2018).

12. Dosztányi, Z., Csizmók, V., Tompa, P. & Simon, I. The pairwise energy content estimated from amino acid composition discriminates between folded and intrinsically unstructured proteins. J. Mol. Biol. 347, 827–839 (2005).

13. Dass, R., Mulder, F. A. A. & Nielsen, J. T. ODiNPred: comprehensive prediction of protein order and disorder. Sci. Rep. 10, 14780 (2020).

14. Hanson, J., Paliwal, K. K., Litfin, T. & Zhou, Y. SPOT-Disorder2: Improved protein intrinsic disorder prediction by ensembled deep learning. Genomics Proteomics Bioinformatics 17, 645–656 (2019).

15. Ishida, T. & Kinoshita, K. PrDOS: prediction of disordered protein regions from amino acid sequence. Nucleic Acids Res. 35, W460–4 (2007).

16. Mizianty, M. J., Peng, Z. & Kurgan, L. MFDp2: Accurate predictor of disorder in proteins by fusion of disorder probabilities, content and profiles. Intrinsically Disord Proteins 1, e24428 (2013).

17. Necci, M., Piovesan, D., CAID Predictors, DisProt Curators & Tosatto, S. C. E. Critical assessment of protein intrinsic disorder prediction. Nat. Methods 18, 472–481 (2021).

18. Conte, A. D., Mehdiabadi, M., Bouhraoua, A., Miguel Monzon, A., Tosatto, S. C. E. & Piovesan, D. Critical assessment of protein intrinsic disorder prediction (CAID) - Results of round 2. Proteins 91, 1925–1934 (2023).

19. Emenecker, R. J., Griffith, D. & Holehouse, A. S. Metapredict: a fast, accurate, and easy-to-use predictor of consensus disorder and structure. Biophys. J. 120, 4312–4319 (2021).

20. Emenecker, R. J., Griffith, D. & Holehouse, A. S. Metapredict V2: An update to metapredict, a fast, accurate, and easy-to-use predictor of consensus disorder and structure. bioRxiv 2022.06.06.494887 (2022) doi:10.1101/2022.06.06.494887.

21. Lotthammer, J. M., Ginell, G. M., Griffith, D., Emenecker, R. J. & Holehouse, A. S. Direct prediction of intrinsically disordered protein conformational properties from sequence. Nat. Methods 21, 465–476 (2024).

22. Mariani, V., Biasini, M., Barbato, A. & Schwede, T. lDDT: a local superposition-free score for comparing protein structures and models using distance difference tests. Bioinformatics 29, 2722–2728 (2013).

23. Jumper, J., Evans, R., Pritzel, A., Green, T., Figurnov, M., Ronneberger, O., Tunyasuvunakool, K., Bates, R., Žídek, A., Potapenko, A., Bridgland, A., Meyer, C., Kohl, S. A. A., Ballard, A. J., Cowie, A., Romera-Paredes, B., Nikolov, S., Jain, R., Adler, J., Back, T., Petersen, S., Reiman, D., Clancy, E., Zielinski, M., Steinegger, M., Pacholska, M., Berghammer, T., Bodenstein, S., Silver, D., Vinyals, O., Senior, A. W., Kavukcuoglu, K., Kohli, P. & Hassabis, D. Highly accurate protein structure prediction with AlphaFold. Nature 596, 583–589 (2021).

24. Alderson, T. R., Pritišanac, I., Kolarić, Đ., Moses, A. M. & Forman-Kay, J. D. Systematic identification of conditionally folded intrinsically disordered regions by AlphaFold2. Proc. Natl. Acad. Sci. U. S. A. 120, e2304302120 (2023).

25. Griffith, D. & Holehouse, A. S. PARROT is a flexible recurrent neural network framework for analysis of large protein datasets. Elife 10, e70576 (2021).

26. Monastyrskyy, B., Fidelis, K., Moult, J., Tramontano, A. & Kryshtafovych, A. Evaluation of disorder predictions in CASP9. Proteins 79 **Suppl 10**, 107–118 (2011).

27. Monastyrskyy, B., Kryshtafovych, A., Moult, J., Tramontano, A. & Fidelis, K. Assessment of protein disorder region predictions in CASP10: Disorder Assessment. Proteins 82 **Suppl 2**, 127–137 (2014).

28. Nielsen, J. T. & Mulder, F. A. A. Quality and bias of protein disorder predictors. Sci. Rep. 9, 5137 (2019).

29. Mehdiabadi, M., Del Conte, A., Nugnes, M. V., Aspromonte, M. C., Tosatto, S. C. E. & Piovesan, D. Critical Assessment of Protein Intrinsic Disorder round 3 - predicting disorder in the era of protein language models. Proteins 94, 414–424 (2026).

30. Aspromonte, M. C., Nugnes, M. V., Quaglia, F., Bouharoua, A., DisProt Consortium, Tosatto, S. C. E. & Piovesan, D.DisProt in 2024: improving function annotation of intrinsically disordered proteins. Nucleic Acids Res. 52, D434–D441 (2024).

31. Berman, H. M., Westbrook, J., Feng, Z., Gilliland, G., Bhat, T. N., Weissig, H., Shindyalov, I. N. & Bourne, P. E. The Protein Data Bank. Nucleic Acids Res. 28, 235–242 (2000).

32. Chicco, D. & Jurman, G. The advantages of the Matthews correlation coefficient (MCC) over F1 score and accuracy in binary classification evaluation. BMC Genomics 21, 6 (2020).

33. Xue, B., Dunker, A. K. & Uversky, V. N. Orderly order in protein intrinsic disorder distribution: disorder in 3500 proteomes from viruses and the three domains of life. J. Biomol. Struct. Dyn. 30, 137–149 (2012).

34. UniProt Consortium. UniProt: The universal protein knowledgebase in 2023. Nucleic Acids Res. 51, D523–D531 (2023).

35. Tretyachenko, V., Vymětal, J., Bednárová, L., Kopecký, V., Jr, Hofbauerová, K., Jindrová, H., Hubálek, M., Souček, R., Konvalinka, J., Vondrášek, J. & Hlouchová, K. Random protein sequences can form defined secondary structures and are well-tolerated in vivo. Sci. Rep. 7, 15449 (2017).

36. Emenecker, R. J., Guadalupe, K., Shamoon, N. M., Sukenik, S. & Holehouse, A. S. Sequence-ensemble-function relationships for disordered proteins in live cells. bioRxivorg 2023.10.29.564547 (2023) doi:10.1101/2023.10.29.564547.

37. Anishchenko, I., Pellock, S. J., Chidyausiku, T. M., Ramelot, T. A., Ovchinnikov, S., Hao, J., Bafna, K., Norn, C., Kang, A., Bera, A. K., DiMaio, F., Carter, L., Chow, C. M., Montelione, G. T. & Baker, D. De novo protein design by deep network hallucination. Nature 600, 547–552 (2021).

38. Pesce, F., Bremer, A., Tesei, G., Hopkins, J. B., Grace, C. R., Mittag, T. & Lindorff-Larsen, K. Design of intrinsically disordered protein variants with diverse structural properties. Sci. Adv. 10, eadm9926 (2024).

39. Yadlowsky, S., Doshi, L. & Tripuraneni, N. Pretraining data mixtures enable narrow model selection capabilities in transformer models. arXiv [cs.LG] (2023) doi:10.48550/arXiv.2311.00871.

40. McCoy, R. T., Yao, S., Friedman, D., Hardy, M. D. & Griffiths, T. L. Embers of autoregression show how large language models are shaped by the problem they are trained to solve. Proc. Natl. Acad. Sci. U. S. A. 121, e2322420121 (2024).

41. Lambert, S. A., Jolma, A., Campitelli, L. F., Das, P. K., Yin, Y., Albu, M., Chen, X., Taipale, J., Hughes, T. R. & Weirauch, M. T. The human transcription factors. Cell 172, 650–665 (2018).

42. Boutet, E., Lieberherr, D., Tognolli, M., Schneider, M. & Bairoch, A. UniProtKB/Swiss-Prot. Methods Mol. Biol. 406, 89–112 (2007).

43. Akiba, T., Sano, S., Yanase, T., Ohta, T. & Koyama, M. Optuna: A next-generation hyperparameter optimization framework. arXiv [cs.LG] (2019).

44. Piovesan, D., Monzon, A. M. & Tosatto, S. C. E. Intrinsic protein disorder and conditional folding in AlphaFoldDB. Protein Sci. 31, e4466 (2022).

45. Tunyasuvunakool, K., Adler, J., Wu, Z., Green, T., Zielinski, M., Žídek, A., Bridgland, A., Cowie, A., Meyer, C., Laydon, A., Velankar, S., Kleywegt, G. J., Bateman, A., Evans, R., Pritzel, A., Figurnov, M., Ronneberger, O., Bates, R., Kohl, S. A. A., Potapenko, A., Ballard, A. J., Romera-Paredes, B., Nikolov, S., Jain, R., Clancy, E., Reiman, D., Petersen, S., Senior, A. W., Kavukcuoglu, K., Birney, E., Kohli, P., Jumper, J. & Hassabis, D. Highly accurate protein structure prediction for the human proteome. Nature 596, 590–596 (2021).

46. Mihara, T., Nishimura, Y., Shimizu, Y., Nishiyama, H., Yoshikawa, G., Uehara, H., Hingamp, P., Goto, S. & Ogata, H. Linking virus genomes with host taxonomy. Viruses 8, 66 (2016).

47. Virtanen, P., Gommers, R., Oliphant, T. E., Haberland, M., Reddy, T., Cournapeau, D., Burovski, E., Peterson, P., Weckesser, W., Bright, J., van der Walt, S. J., Brett, M., Wilson, J., Millman, K. J., Mayorov, N., Nelson, A. R. J., Jones, E., Kern, R., Larson, E., Carey, C. J., Polat, İ., Feng, Y., Moore, E. W., VanderPlas, J., Laxalde, D., Perktold, J., Cimrman, R., Henriksen, I., Quintero, E. A., Harris, C. R., Archibald, A. M., Ribeiro, A. H., Pedregosa, F., van Mulbregt, P. & SciPy 1.0 Contributors. SciPy 1.0: fundamental algorithms for scientific computing in Python. Nat. Methods 17, 261–272 (2020).

48. Chamberlain, S. A. & Szöcs, E. taxize: taxonomic search and retrieval in R. F1000Res. 2, 191 (2013).

49. Letunic, I. & Bork, P. Interactive Tree of Life (iTOL) v6: recent updates to the phylogenetic tree display and annotation tool. Nucleic Acids Res. 52, W78–W82 (2024).

50. Mao, A. H., Crick, S. L., Vitalis, A., Chicoine, C. L. & Pappu, R. V. Net charge per residue modulates conformational ensembles of intrinsically disordered proteins. Proc. Natl. Acad. Sci. U. S. A. 107, 8183–8188 (2010).

51. Das, R. K. & Pappu, R. V. Conformations of intrinsically disordered proteins are influenced by linear sequence distributions of oppositely charged residues. Proc. Natl. Acad. Sci. U. S. A. 110, 13392–13397 (2013).

52. Kyte, J. & Doolittle, R. F. A simple method for displaying the hydropathic character of a protein. J. Mol. Biol. 157, 105–132 (1982).

53. Holehouse, A. S., Das, R. K., Ahad, J. N., Richardson, M. O. G. & Pappu, R. V. CIDER: Resources to analyze sequence-ensemble relationships of intrinsically disordered proteins. Biophys. J. 112, 16–21 (2017).

54. Alston, J. J., Ginell, G. M., Soranno, A. & Holehouse, A. S. The analytical Flory random coil is a simple-to-use reference model for unfolded and disordered proteins. J. Phys. Chem. B 127, 4746–4760 (2023).

