## Supplementary Information for "Metapredict enables accurate disorder prediction across the Tree of Life"

### SUPPLEMENTARY INFORMATION FOR:
#### Metapredict enables accurate disorder prediction across the Tree of Life

Jeffrey M. Lotthammer[1, 2], Jorge Hernández-García[3], Daniel Griffith[1, 2], Dolf Weijers[3], Alex S. Holehouse[1, 2, 4], Ryan J. Emenecker[1,2,*]

[1]Department of Biochemistry and Molecular Biophysics, Washington University School of Medicine, St. Louis, MO, 63100, USA
[2]Center for Biomolecular Condensates, Washington University, St. Louis, MO 63130, USA
[3]Centro de Biotecnologıa y Genomica de Plantas, Universidad Politecnica de Madrid (UPM) – Instituto Nacional de Investigacion y Tecnologıa, Agraria y Alimentaria (INIA/CSIC), Campus de Montegancedo UPM, Pozuelo de Alarcon, 28223, Spain
[4]Laboratory of Biochemistry, Wageningen University, Stippeneng 4, 6708WE Wageningen, The Netherlands

[5]Lead contact

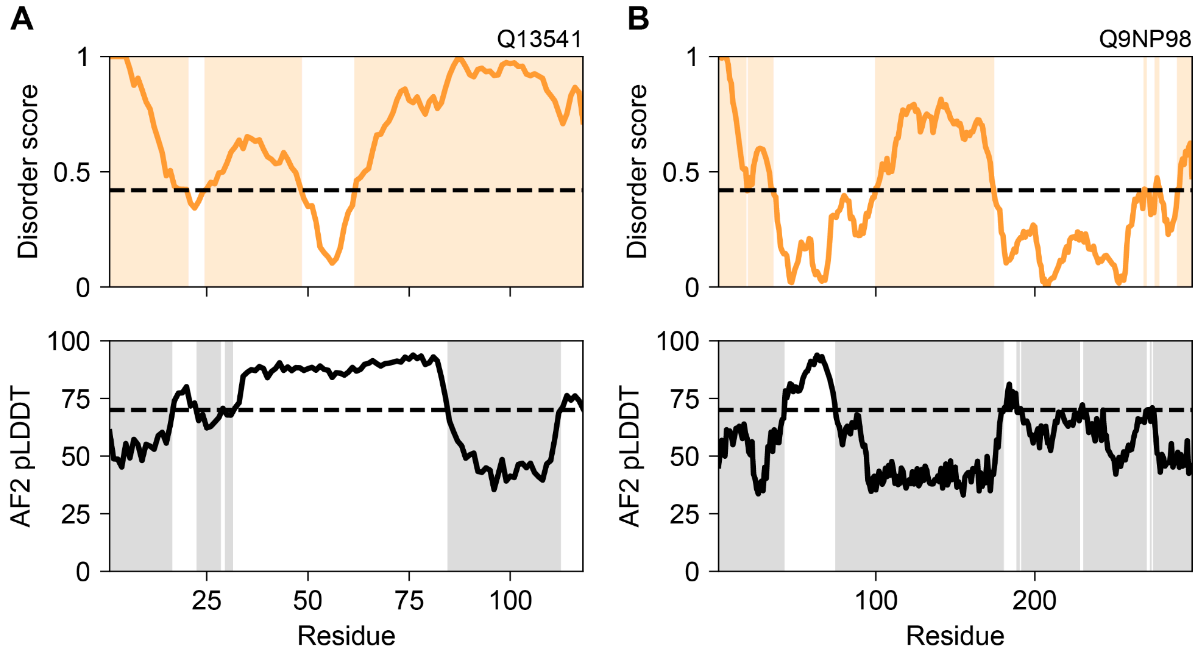


**Supplemental Fig. 1**. **AF2 pLDDT scores and metapredict V1 disorder scores identify different protein regions as disordered.** Examples of how pLDDT and metapredict V1 scores differ across different sequences. **(A)** Disorder scores were generated using the metapredict V1 network (top) and AF2 pLDDT values (bottom) for *Homo sapiens* EIF4EBP1 (UniProt Q13541). For disorder scores (top), highlighted regions are above the 0.42 threshold and are thus classified as disordered. For AF2 pLDDT values (bottom), scores below 70 (classified as low or very low confidence by AF2) are highlighted. **(B)** Same as for (A) except for *Homo sapiens* Myozenin-1 (UniProt Q9NP98).

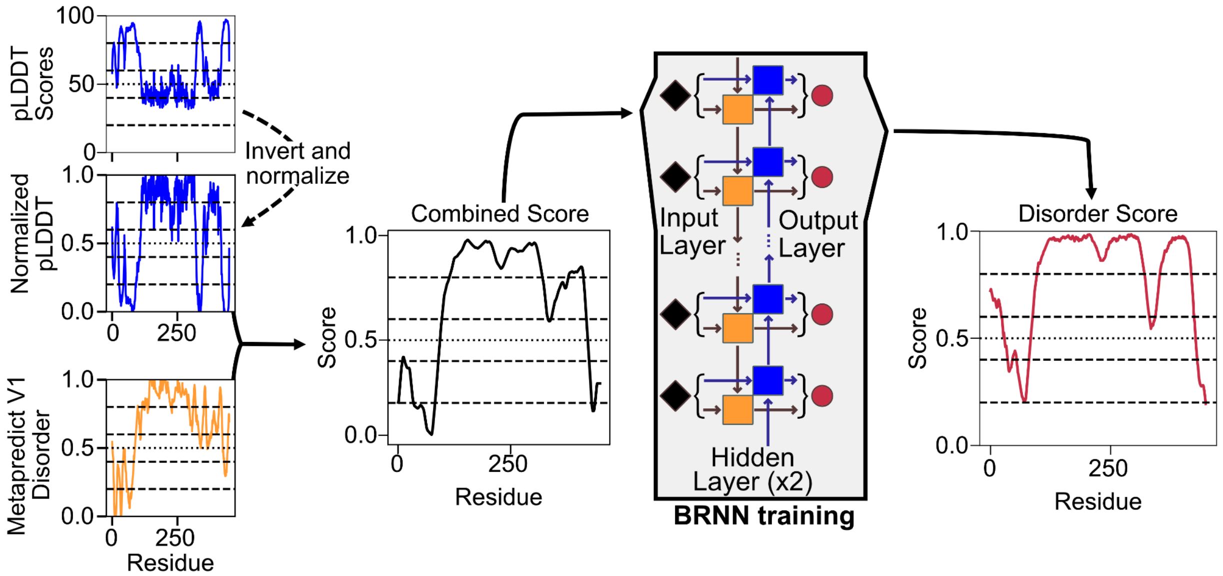


**Supplemental Fig. 2. Generating combined disorder scores from metapredict V1 disorder scores and AF2 pLDDT.** The approach for generating the 'combined disorder scores' for metapredict V3 was as follows: ***(1)*** pLDDT scores (top left) were divided by 100 and then were normalized using a base value of 0.35 and a ceiling value of 0.95, meaning final values lie between 0 and 1. We inverted these scores so that 1 = fully disordered and 0 = fully ordered (left side, middle). This generated "re-cast" pLDDT scores; ***(2)*** metapredict V1 disorder scores (left side, bottom) were combined with our re-cast pLDDT scores. If either value was greater than 0.5, we used the higher of the two; if both were below 0.5, we used the lower of the two. We then applied a Savitzky-Golay smoothing filter to these scores, clipped scores above 1 to 1, and clipped scores below 0 to 0. Finally, we applied a moving-average smoothing step with a window size of 25. We then generated these combined scores (graph labeled 'combined') for 455,666 protein sequences and used them to train a BRNN-LSTM. The scores generated by the resultant network (furthest right graph) accurately recapitulate the 'combined disorder scores.'

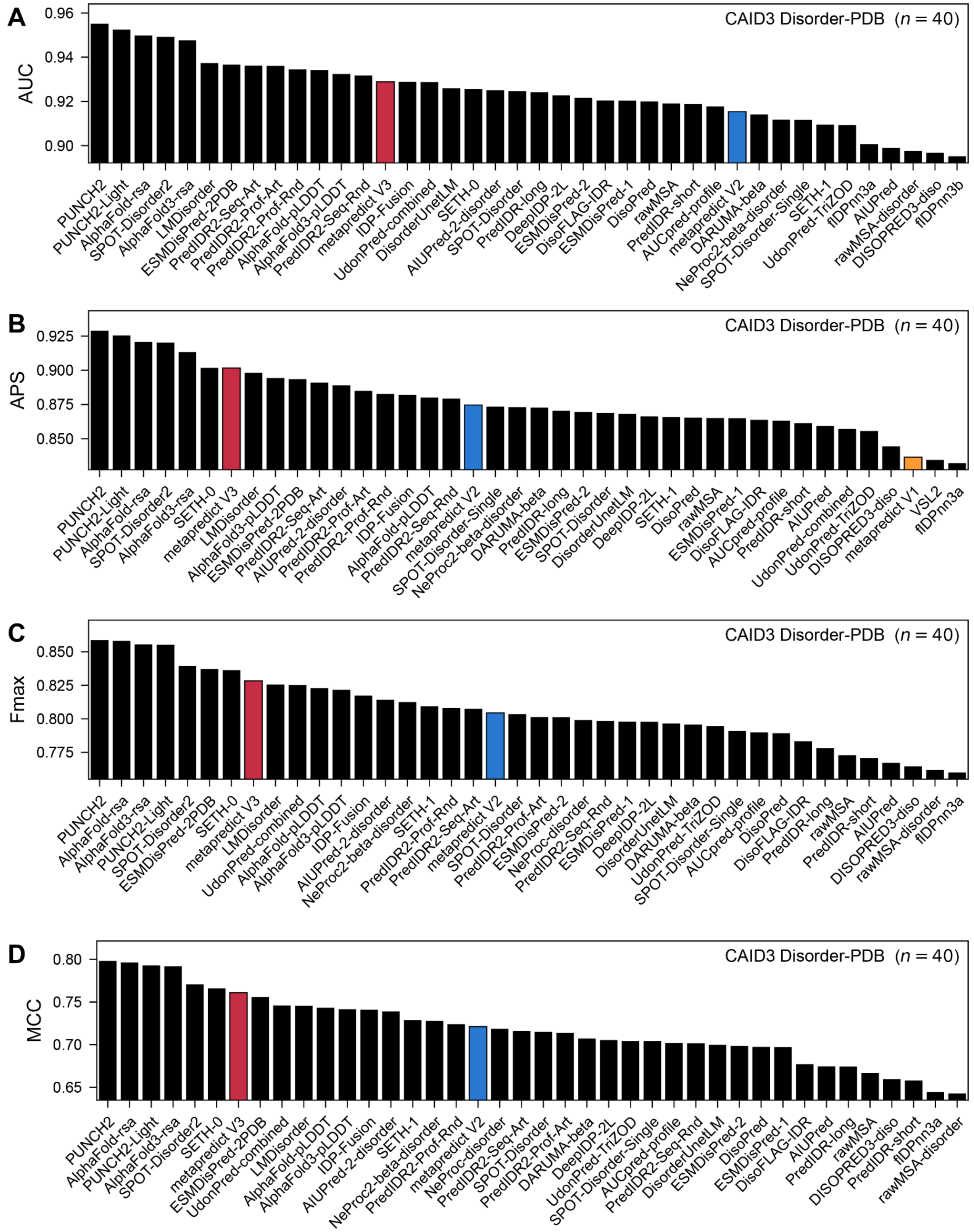


**Supplemental Fig. 3. CAID3 rankings for metapredict 1, 2 and 3 across different evaluation criteria.** All 66 CAID3 disorder predictors on the CAID3 Disorder-PDB dataset (319 proteins). For the Disorder-PDB dataset, residues annotated as disordered in DisProt count as disordered (31,401), and residues resolved in the PDB count as ordered (67,838); all other residues are excluded. Each panel shows the top 40 predictors for one metric, with metapredict V1 (orange), V2 (blue), and V3 (crimson) coloured where they appear. Across the four metrics, metapredict V3 ranks 14th, 7th, 8th and 7th and V2 ranks 18th-30th. V1 ranks 38th-45th, so it appears only in (B). **(A)** AUC, the area under the ROC curve: the probability that a randomly chosen disordered residue scores higher than a randomly chosen ordered one (1 is perfect, 0.5 is random). **(B)** APS (average precision), the area under the precision-recall curve. When disordered residues are the minority it is a stricter test than AUC, because ordered residues among the top scores pull it down. A random predictor scores 0.32, the fraction of residues that are disordered. **(C)** Fmax, the highest F1 score (the harmonic mean of precision and recall) over all score cutoffs. **(D)** MCC (Matthews correlation coefficient) at each predictor's Fmax cutoff, from −1 (always wrong) through 0 (random) to 1 (perfect). This MCC uses each predictor's best cutoff, not its own yes/no calls, so it differs from the MCC in Figure 1C.

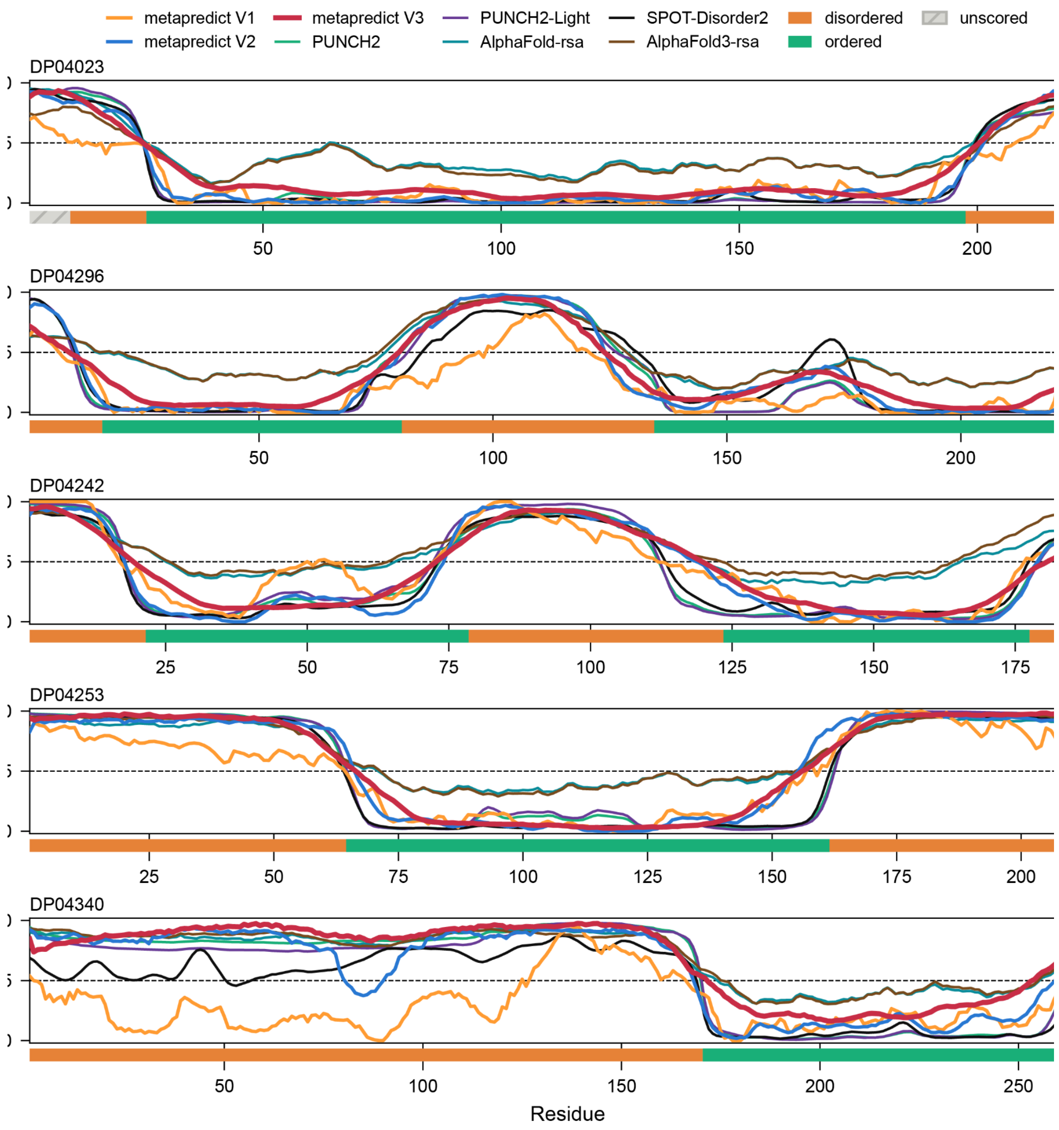


**Supplemental Fig. 4. Disorder profiles from metapredict V1, V2, and V3 and the top CAID3 predictors for five CAID3 Disorder-PDB proteins.** Per-residue disorder scores from metapredict V1 (orange), V2 (blue), and V3 (crimson), and the five CAID3 disorder predictors with the highest AUC (PUNCH2, PUNCH2-Light, AlphaFold-rsa, SPOT-Disorder2, and AlphaFold3-rsa). Red shading highlights regions Disorder-PDB categorizes as disordered. Unshaded residues are ordered, except residues 1–9 of DP04023, which are excluded from scoring.

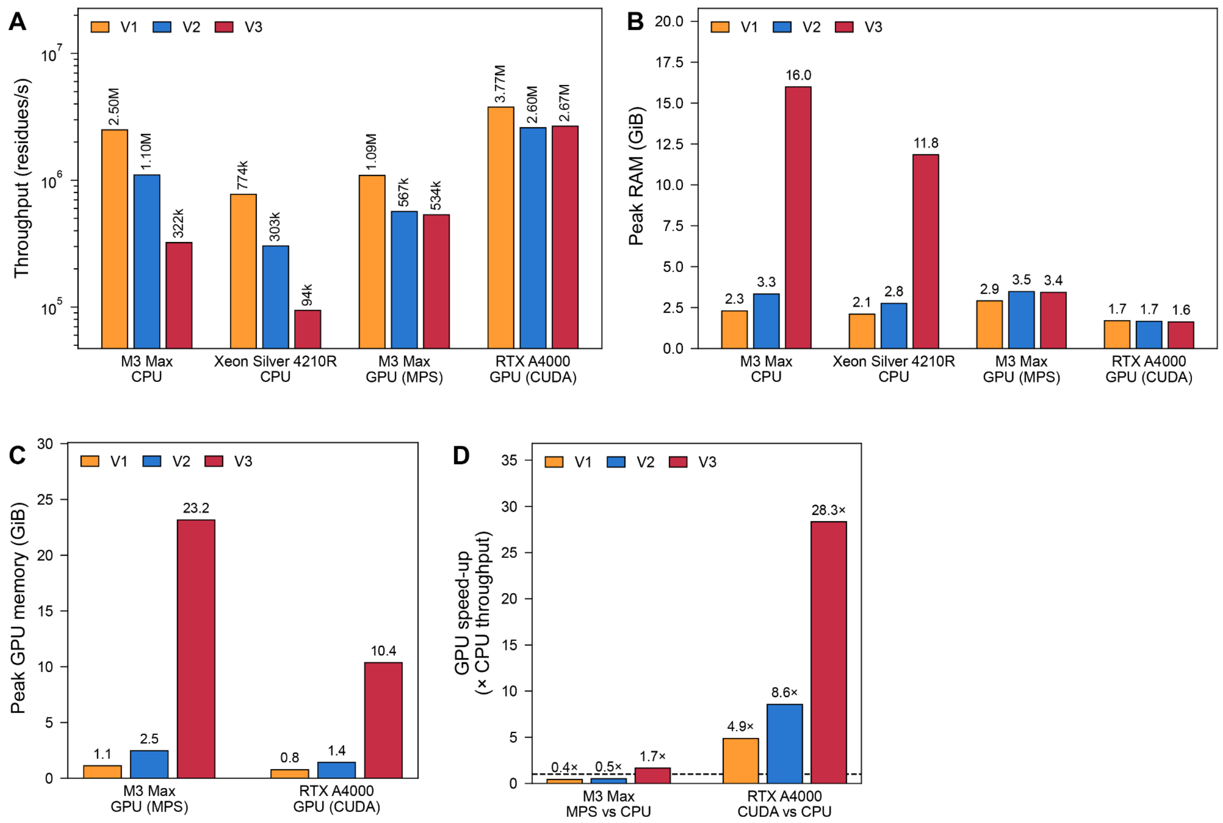


**Supplemental Fig. 5. Prediction speed and memory use of metapredict V1, V2, and V3 on CPUs and GPUs.** We benchmarked Metapredict using the UniProt human reference proteome (UP000005640; 20,652 sequences, 11,453,425 residues). Runs were performed using metapredict's device-specific default batch size (256 on CPU and CUDA, 512 on MPS). **(A)** Throughput (residues predicted per second). **(B)** Peak resident memory of the Python process during prediction. Peak memory is largely determined by the longest sequence (Titin; 34K residues), and truncating sequence lengths substantially reduces the maximum memory footprint. **(C)** Peak GPU memory during prediction. For CUDA, this is the maximum memory PyTorch allocates; for MPS, it is the largest sampled Metal driver allocation. On the M3 Max, the GPU shares the 128 GB of system memory. As with CPU-relevant memory, peak memory largely depends on the longest sequence. **(D)** GPU speed-up: GPU throughput divided by CPU throughput. On Apple Silicon, CPU implementation for V1 and V2 is faster than MPS, whereas where CUDA-compatible GPUs are available, GPUs offer substantial improvements over CPU performance.

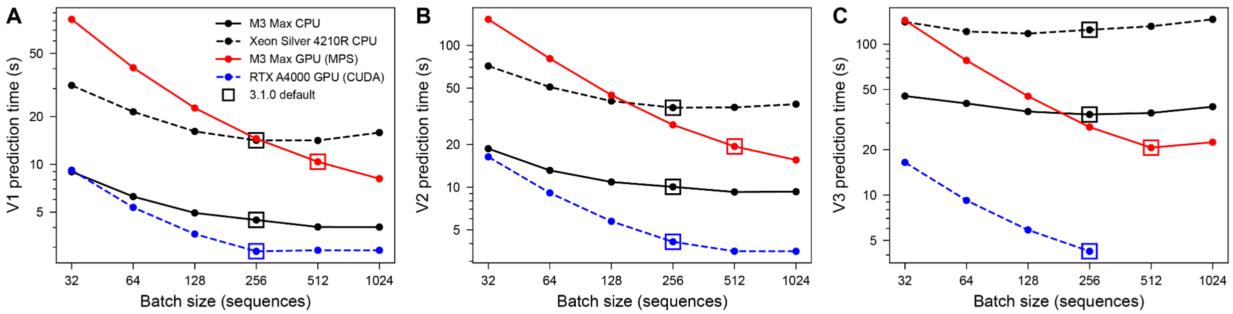


**Supplemental Fig. 6. Batch-size dependence of prediction time.** Time to predict the human proteome (20,652 sequences) with metapredict **(A)** V1, **(B)** V2, and **(C)** V3 at batch sizes from 32 to 1,024. Open squares mark the default batch size for each network on for each device. CPU points are the median of three sweeps; GPU points are single runs. V3 ran out of the RTX A4000's 16 GB of memory at batch sizes of 512 and 1,024, so its CUDA line stops at 256. We chose default batch sizes to balance performance while trading off memory footprint. Batch size can be set manually as needed.

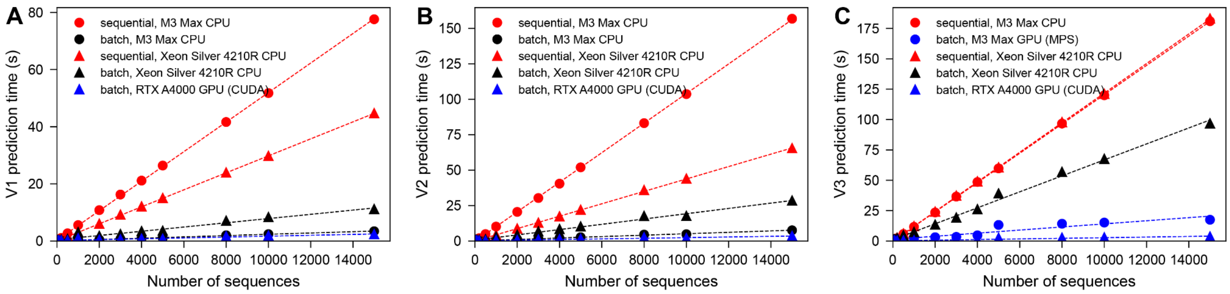


**Supplemental Fig. 7. Batch vs. sequential prediction performance.** Time to predict random subsets of 10 to 15,000 human proteins with metapredict **(A)** V1, **(B)** V2, and **(C)** V3. Sequences were predicted either one per call (sequential, red, always on the CPU) or in one batched call using each network's default batch size, on its default device on the MacBook Pro (CPU for V1 and V2, MPS for V3) and on both the CPU and the GPU on the workstation. Batch prediction accords substantial improvements in predicting across large sets of sequences.

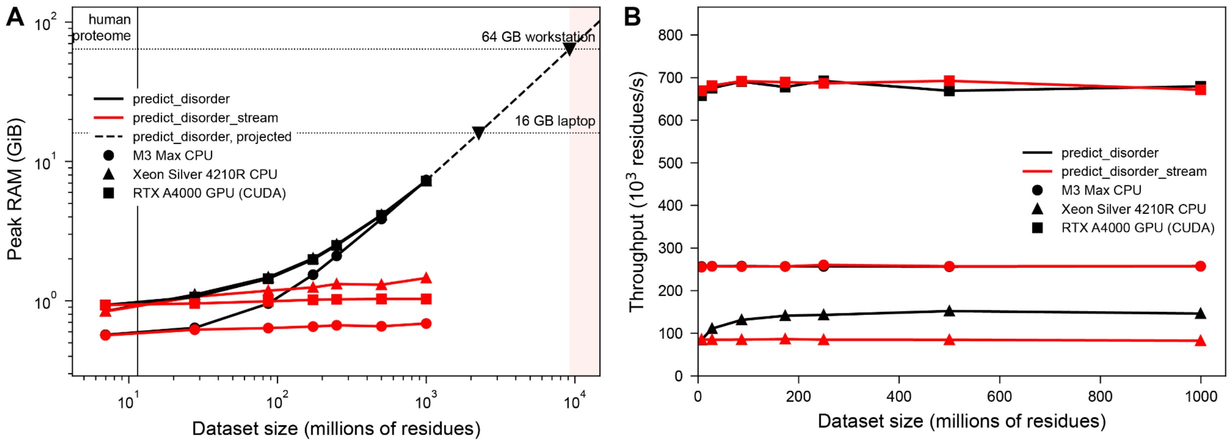


**Supplemental Fig. 8. Streaming prediction keeps memory bounded as datasets grow.** We generated synthetic datasets of 20,000 to 2,880,000 random sequences (7.0 million to 1.0 billion residues; lengths log-normally distributed with a median of ~300 residues and limited to 40–2,500) and predicted them with metapredict V3. We used either **`predict_disorder`** (black), which holds the whole dataset and all predictions in memory, or **`predict_disorder_stream`** (red), which reads, predicts, and returns 20,000 sequences at a time (the metapredict 3.1.0 default chunk size). Both used a batch size of 32. Each point was measured in a fresh process on an Apple M3 Max CPU (circles), an Intel Xeon Silver 4210R CPU (triangles), or an NVIDIA RTX A4000 GPU (squares). **(A)** Peak RAM versus dataset size; for the GPU, this is host RAM only. **`predict_disorder`** memory grows linearly with dataset size on all three devices, reaching 7.2–7.4 GiB at 1 billion residues. Extrapolating the M3 Max points (dashed line) predicts that predict_disorder would fill a 16 GB laptop at 2.25 billion residues and a 64 GB workstation at 9.2 billion (inverted triangles). **`predict_disorder_stream`** used at most 1.5 GiB at every size. Its memory is flat on the M3 Max (0.57–0.69 GB) and the A4000 (0.93–1.03 GB) but creeps up slowly on the Xeon CPU (0.84 to 1.46 GB). This is because PyTorch's oneDNN library caches LSTM kernels for each new batch shape (up to a fixed number), and the Linux (glibc) memory allocator holds on to freed memory. With oneDNN's cache turned off and the allocator set to return large blocks immediately, streaming memory on the Xeon stayed flat in a 200,000-sequence test. The vertical line marks the size of the human proteome (11.5 million residues). **(B)** Throughput versus dataset size. On the A4000 and the M3 Max, both functions run at the same speed at every size (660–690 and ~257 thousand residues/s, respectively), so streaming carries no speed cost there. On the Xeon CPU, **`predict_disorder_stream`** runs at a constant ~85 thousand residues/s, while **`predict_disorder`** speeds up from 85 to ~150 thousand residues/s as the dataset grows. This reflects how PyTorch runs LSTMs on x86 CPUs: a batch goes to oneDNN's fast LSTM kernel only if every sequence has the same length; otherwise, it uses a slower generic implementation. metapredict sorts sequences by length before batching, so the larger the dataset, the more sequences share each length and the more batches contain a single length (16% of residues at 20,000 sequences, 96% at 2.88 million).

**predict_disorder_stream** sorts each 20,000-sequence chunk on its own, so this fraction stays at ~16% whatever the dataset size, and its throughput stays at that of predict_disorder on 20,000 sequences. Neither the M3 Max nor the GPU uses this oneDNN path, so their throughput does not depend on batch composition.

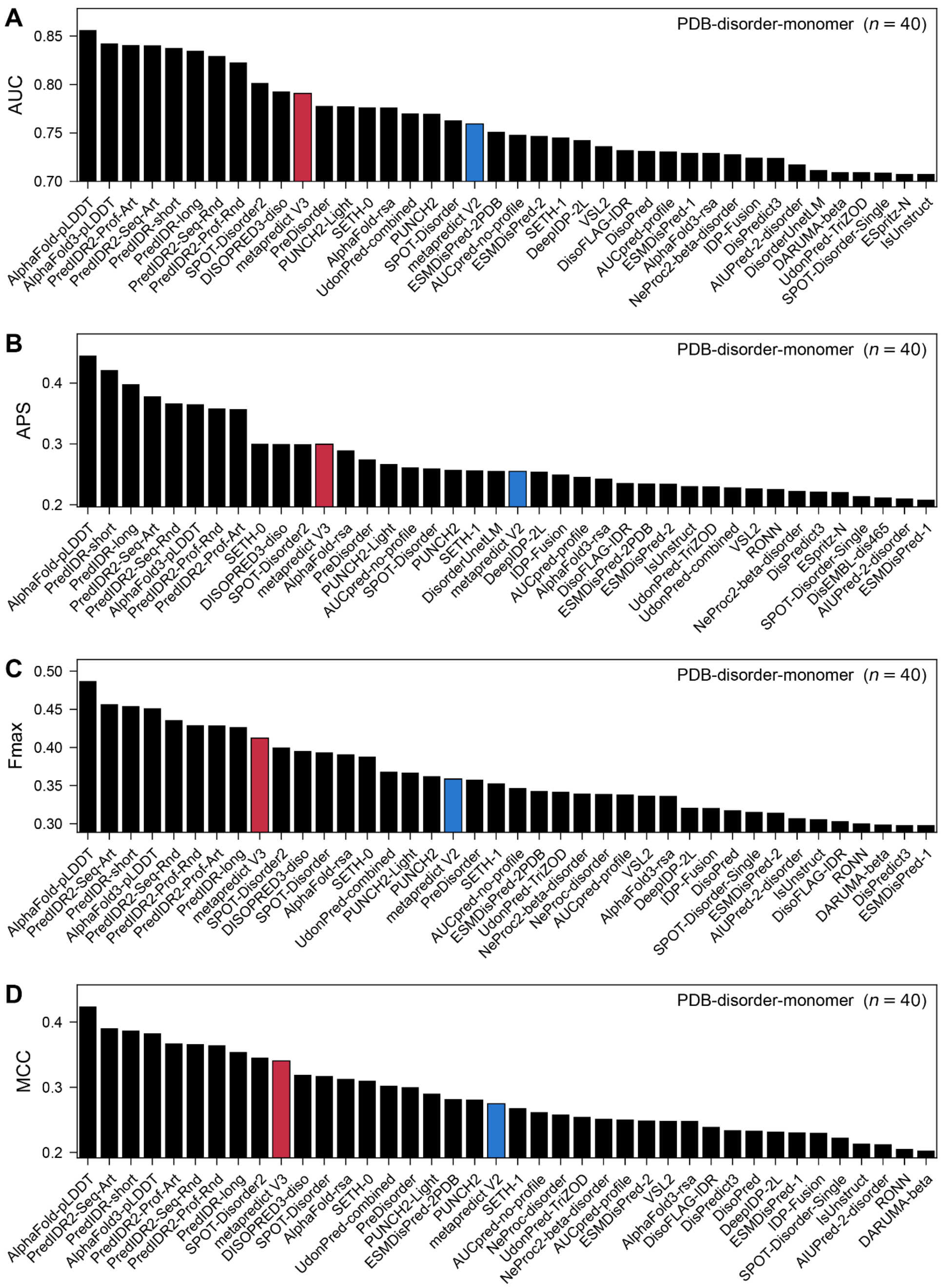


**Supplemental Fig. 9. CAID3 predictor rankings on a stringent PDB-disorder-monomer benchmark.** This benchmark uses only PDB structures of isolated, monomeric proteins. A residue is ordered if it is resolved in such a structure, and disordered if it lies within the construct but has no electron density. All other residues are excluded, leaving 134 proteins with 32,904 ordered and 3,807 disordered residues (down from 319 proteins in the full PDB-disorder set). The panels rank the top 40 of 66 CAID3 disorder predictors by **(A)** AUC, **(B)** APS, **(C)** Fmax, and **(D)** MCC at each predictor's Fmax threshold, as described in **Supplemental Fig. S3**. Metapredict V1 (orange), V2 (blue), and V3 (crimson). Metapredict V3 ranks 11th, 12th, 9th, and 10th on the four metrics, and AlphaFold-pLDDT ranks first on all four. We note this analysis is somewhat flawed, as much of the signal comes from a single entry (DP04052), which is too long for some methods. As such, we advise against reading into this analysis - we performed it to investigate if/how it changed any of the rankings, but did not see any evidence that disorder prediction (in general) was biased by intermolecular interactions in the structural data.

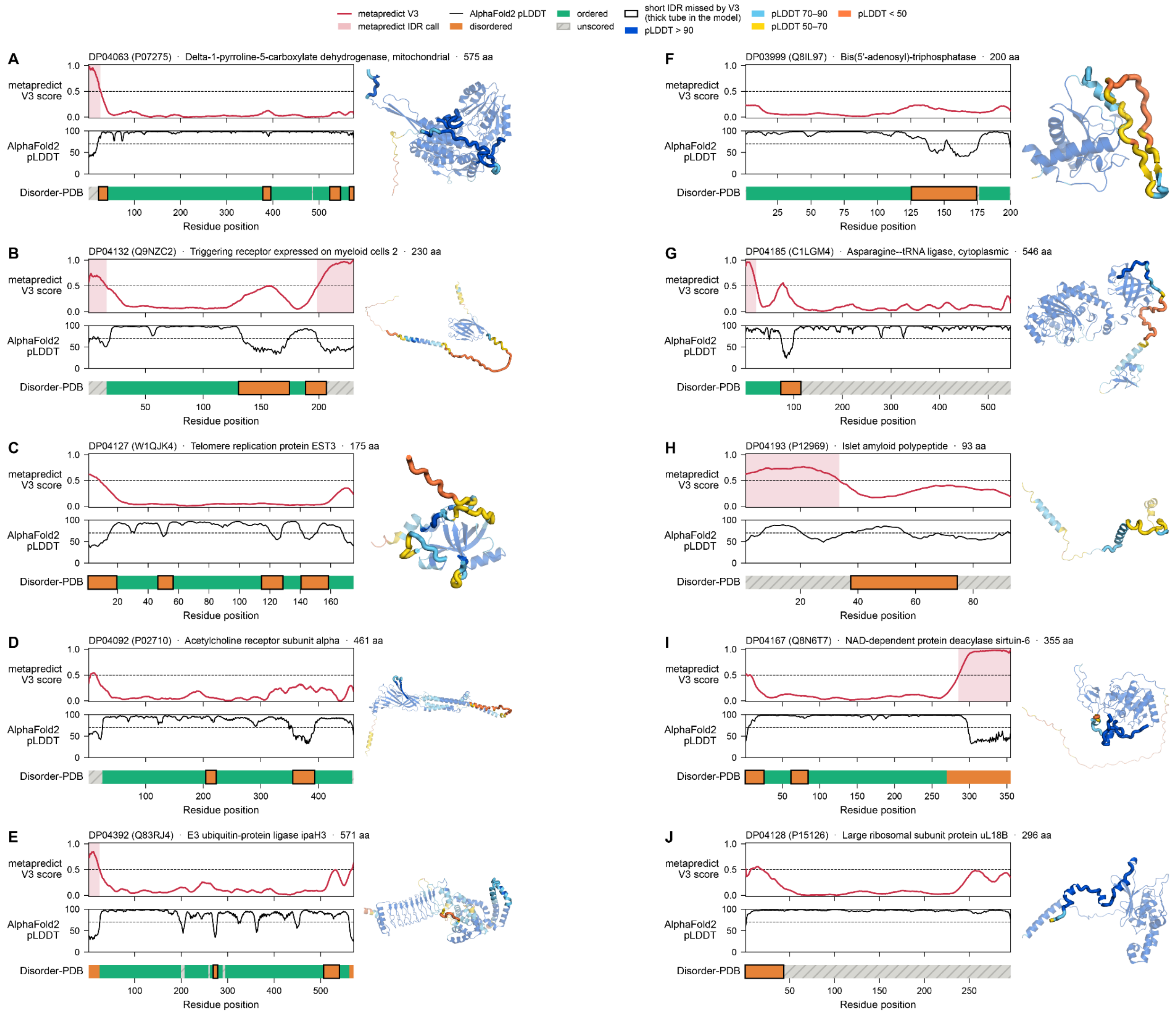


**Supplemental Fig. 10. Examples of short IDRs that metapredict V3 misses.** Ten Disorder-PDB proteins with short IDRs (10–49 residues) that metapredict V3 misses. V3 misses an IDR when fewer than half of its residues score at least 0.5 and fewer than half lie in a predicted IDR. Examples highlight three failure modes: **(A-E)** Loops (IDRs between ordered residues); **(F-H)** IDRs adjacent to unscored residues (i.e., residues not annotated as disordered or resolved in the PDB); and **(I–J)** IDRs at the N- or C-terminus of proteins. Each panel shows three tracks: (1) The metapredict V3 score, with any predicted IDRs shaded in red, (2) the AlphaFold2 pLDDT scores, from CAID3's AlphaFold-pLDDT predictions; and (3) the Disorder-PDB labels, with the missed IDRs outlined in thick black lines. For each protein, we also provide the AlphaFold2-predicted structure colored by pLDDT with the regions missed highlighted as thicker tube representations.

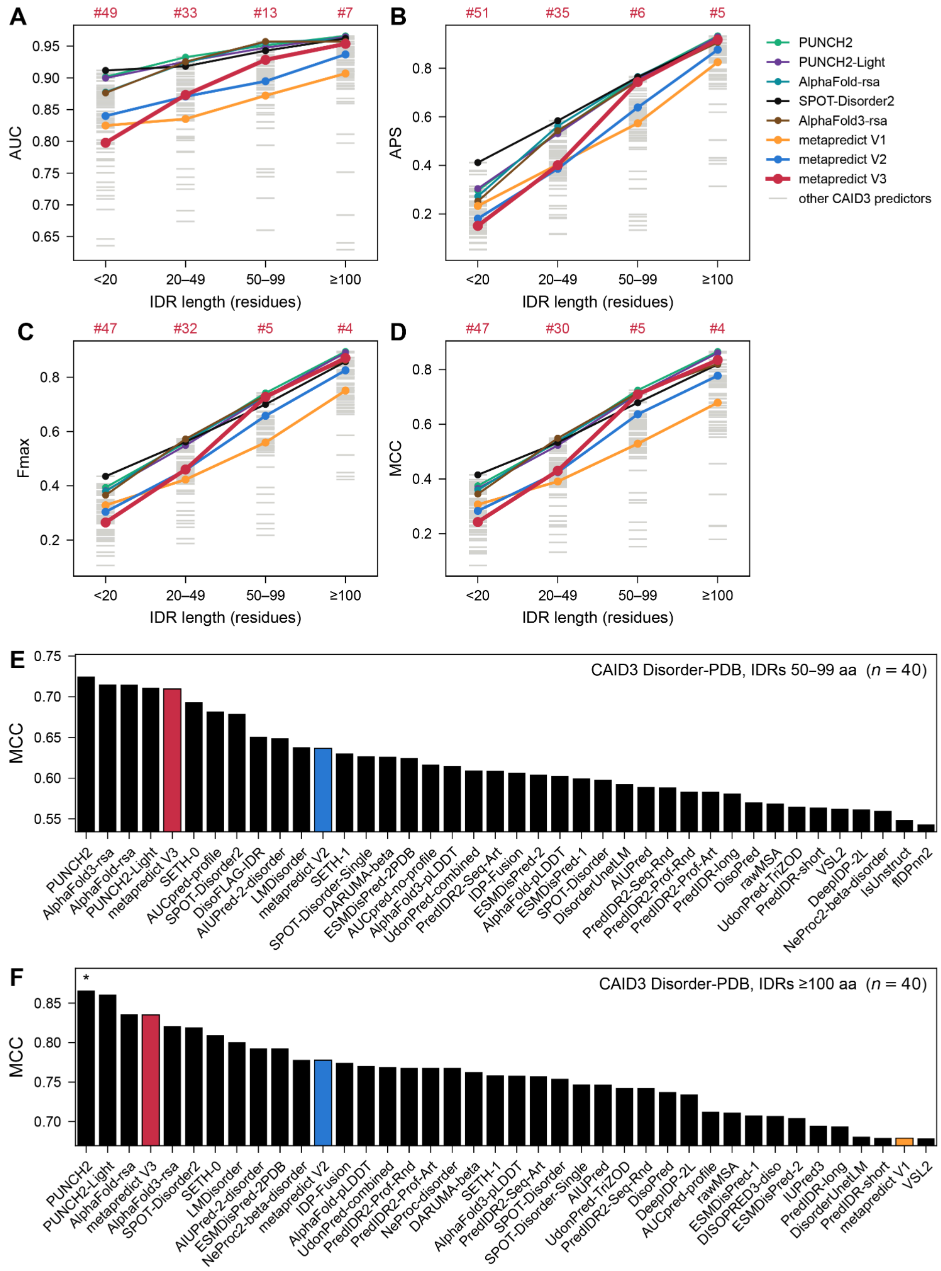


**Supplemental Fig. 11. Metapredict V3 performs best for long IDRs**. CAID3 Disorder-PDB scored separately by IDR length. Bins are: 10-19 residues (n=158 IDRs), 20-49 residues (n=135 IDRs), 50-99 residues (n=68 IDRs), and 100 or more residues (n=94 IDRs). For each group, we calculated **(A)** AUC, **(B)** APS, **(C)** Fmax, and **(D)** MCC at each predictor's Fmax threshold for all 66 CAID3 disorder predictors (grey) in each bin. The five top CAID3 predictors and metapredict V1, V2, and V3 are drawn as thick colored lines, the remaining predictors as thin grey lines, and the number above each bin is metapredict V3's rank. Metapredict V3 is noticeably poor at short IDRs, but highly accurate for long (≥50 residue) IDRs. Broadly speaking, metapredict V3 does not do well at identifying disordered loops or short N/C-terminal tails adjacent to folded domains, but is highly accurate for identifying larger, "*bona fide*" disordered domains. **(E)** The top 40 predictors by MCC for IDRs of 50–99 residues. An asterisk marks a predictor significantly more accurate than metapredict V3 (paired bootstrap over proteins, 1,000 resamples, 95% interval). No predictors are statistically more accurate than metapredict V3 in this regime. **(F)** The top 40 predictors by MCC for IDRs that are 100 residues or longer. An asterisk marks a predictor significantly more accurate than metapredict V3 (paired bootstrap over proteins, 1,000 resamples, 95% interval). Only PUNCH2 is statistically more accurate than metapredict V3 in this regime. Overall, our analysis suggests that for IDRs 50 residues or longer, only PUNCH2 is statistically more accurate than metapredict V3.

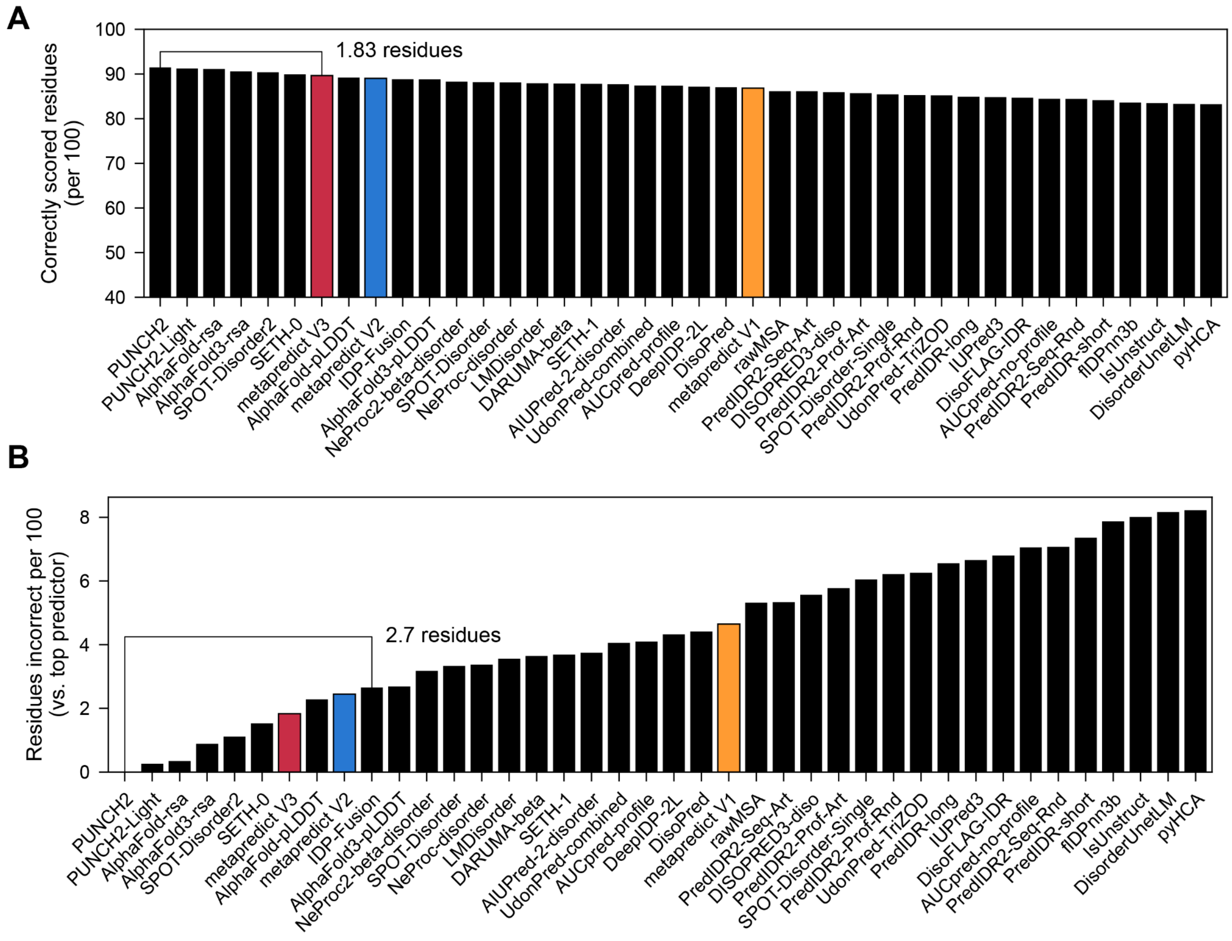


**Supplemental Fig. 12. The top CAID3 disorder predictors differ by only a few residues per 100. (A)** Correctly scored residues per 100 on the CAID3 Disorder-PDB dataset, for the 40 most accurate CAID3 predictors (of 66). Residues are pooled across all 319 proteins. Each predictor is scored on its submitted binary calls where available, or otherwise with a cutoff of 0.5 (0.581 and 0.312 for the AlphaFold2 and AlphaFold3 RSA and pLDDT tracks, the published AlphaFold2 cutoffs). Metapredict is scored on residues that fall within DisorderedDomains. Inclusion and cutoffs are as in **Fig. 1C**. Six of the 40 predictors shown are cut at 0.5 (LMDisorder, AIUPred-2-disorder, UdonPred-combined, rawMSA, UdonPred-TriZOD and DisoFLAG-IDR). The bracket shows the gap between the best predictor (PUNCH2, 91.5) and metapredict V3 (89.6, 7th), which is 1.8 residues per 100. **(B)** Analogous analysis for extra residues scored incorrectly per 100, relative to the best predictor. The bracket shows the gap between the best and the tenth-best predictor: 2.7 residues per 100.

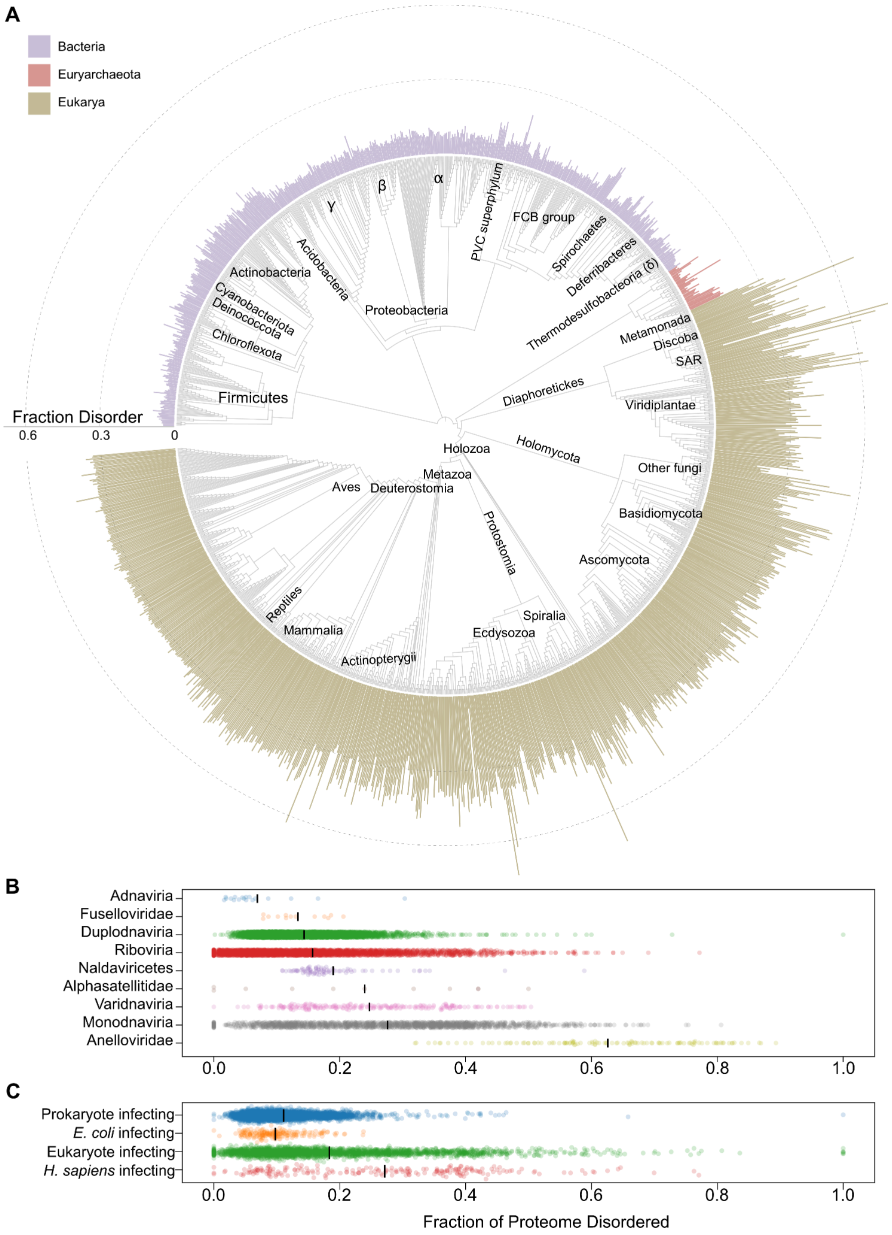


**Supplemental Fig. 13. Disorder across the tree of life, quantified as the total fraction across the proteome**. Re-plotting data from **Fig. 3** with each proteome's disorder fraction calculated as the number of residues predicted to be disordered divided by its total number of residues (total fraction across proteome), rather than the mean fraction disordered per protein (used in **Fig. 3**). This measure gives more weight to long proteins. **(A)** Proteome-wide disorder across 1,480 families (12,568 species), with major taxonomic lineages labeled. **(B)** Viral proteomes grouped by lineage (only lineages with ≥10 proteomes; single-gene proteomes excluded). **(C)** Viral proteomes grouped by host. In (B) and (C), black lines mark the mean.

Viruses that infect eukaryotes are more disordered than viruses that infect prokaryotes (KS test, $p = 1 \times 10^{-122}$). Likewise, human-infecting viruses are more disordered than E. coli-infecting viruses (KS test, $p = 1 \times 10^{-36}$).

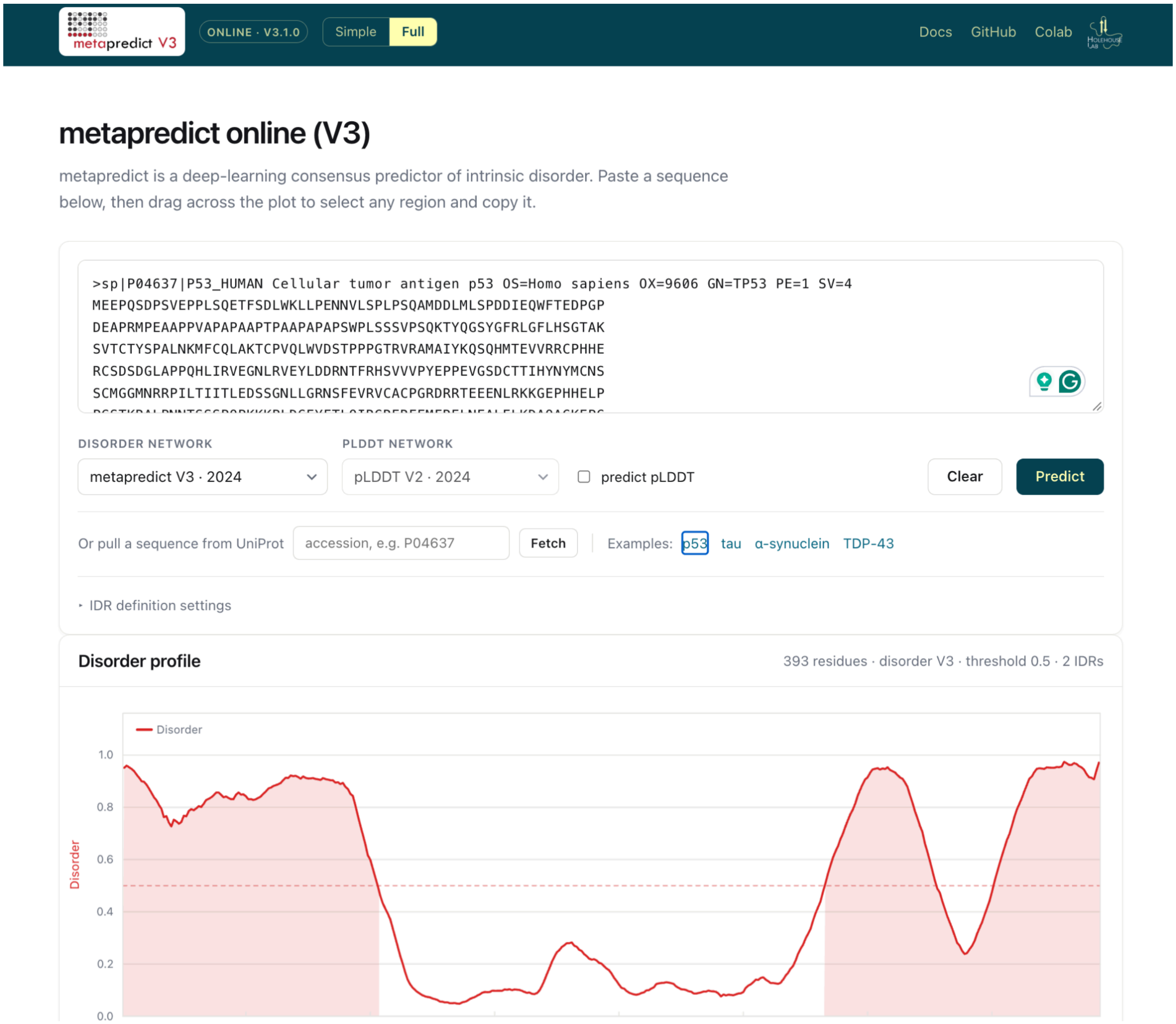


**Supplemental Fig. 14.** Screenshot of the metapredict online web server. The V3 web server offers substantial feature upgrades over our previous V1 server, including the ability to select different predictors, customize algorithmic decisions for IDR calling, and calculate sequence properties across different IDRs.

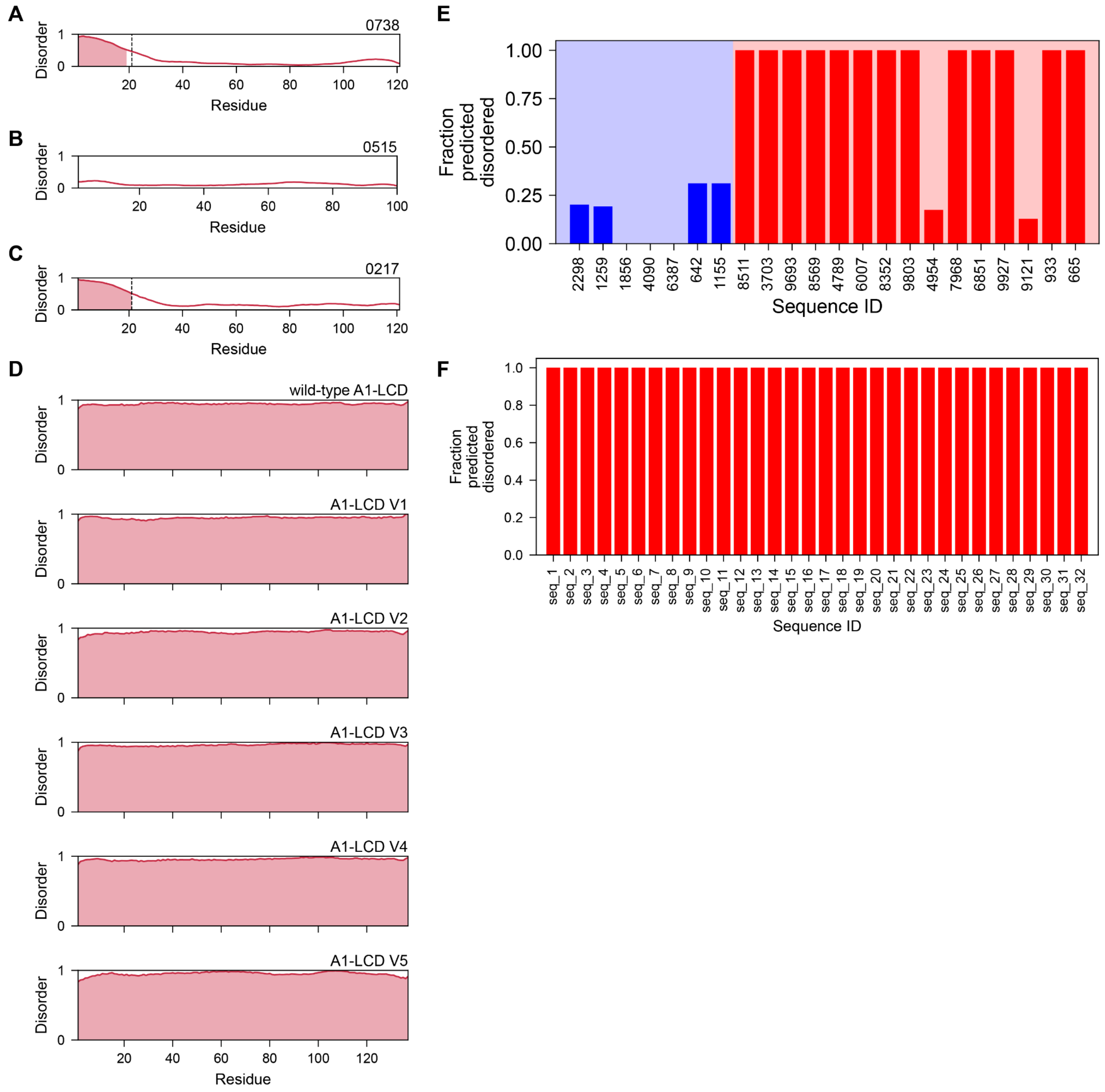


**Supplemental Fig. 15. Evaluation of metapredict V3 on synthetic proteins. (A, B, and C)** Artificial folded proteins are designed through deep hallucination[1]. In sequences (A) and (C), an N-terminal HIS tag is present; metapredict V3 correctly predicts that the HIS tag will be disordered and the remainder of the protein will be folded. **(D)** Pesce et al.[2] designed and tested synthetic variants of the hnRNPA1 low complexity domain *in vitro* (wild-type at top). All variants are experimentally verified as being disordered. **(E)** Random peptide sequences assessed *in vitro* by Tretyachenko *et al.*, folded (left-hand seven sequences in the blue shaded region), and disordered (right-hand 15 sequences in the red shaded region), were assessed[3]. **(F)** Hunter *et al.*[4] designed 32 synthetic disordered proteins. All sequences were tested in cells and found to be disordered based on inferred end-to-end distance.

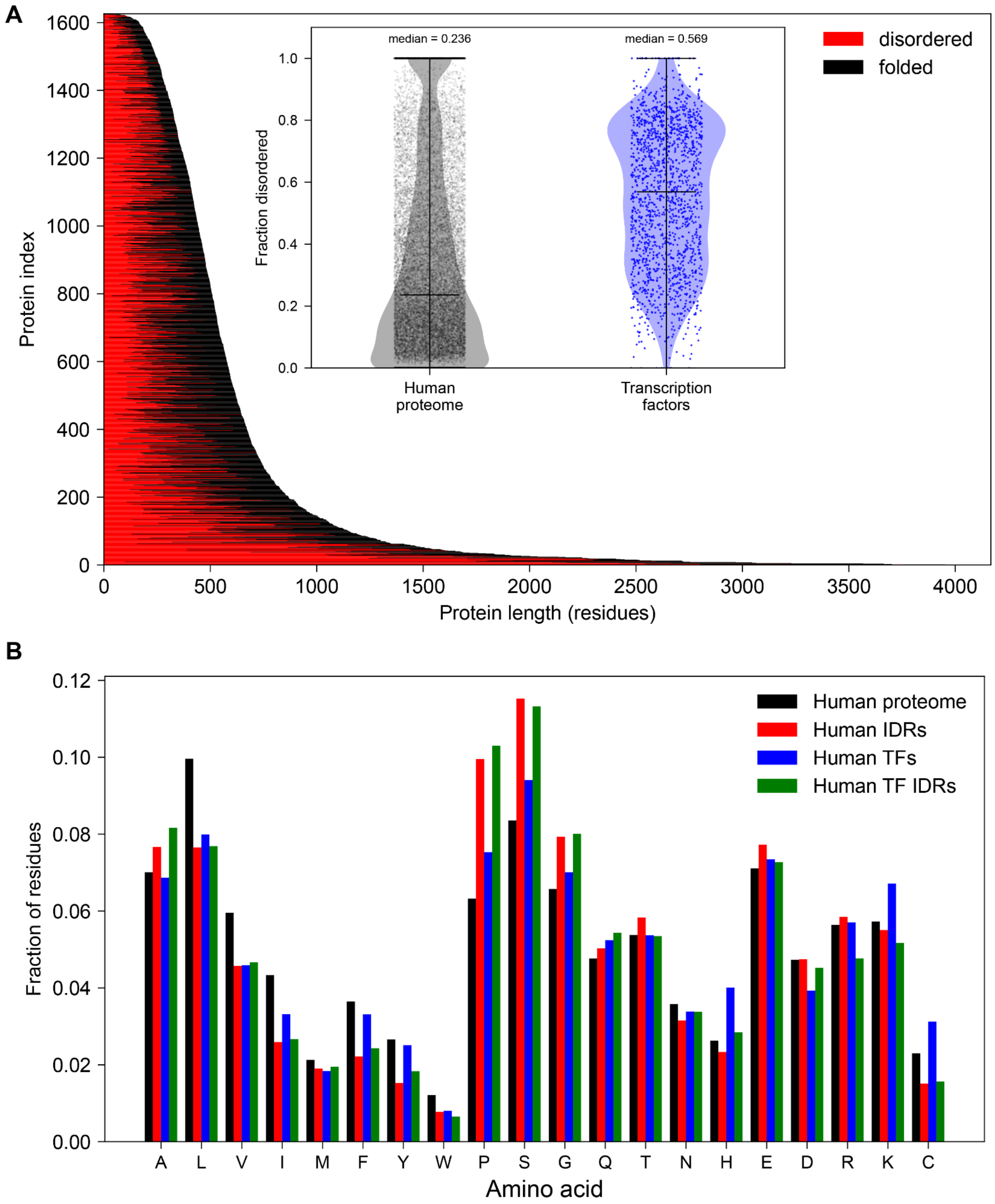


**Supplemental Fig. 16. Analysis of human transcription factors.**

**(A)** 1608 human transcription factors are represented as a stacked bar plot ordered by total protein length[5]. For each protein (indexed along the y-axis), red bars show the number of residues within IDRs, while black bars show the number of residues in folded domains. The inset compares human transcription factors vs. the human proteome; human transcription factors are (by median) ~2.4x more disordered than proteins in the human proteome. The inset quantifies disorder as the fraction of residues in disordered regions. **(B)** Comparison of amino acid composition in [from left-to-right] the human proteome (black), all human IDRs (red), all human transcription factors (blue), and human transcription factor IDRs (green). Human transcription factors show enrichment for lysine, cysteine, and histidine (found in DNA-binding domains), but overall, human transcription factor IDRs and all human proteome IDRs are compositionally quite similar.

#### SUPPLEMENTARY REFERENCES

10.1038/s41592-021-01117-3

1. Anishchenko I, Pellock S J, Chidyausiku T M, Ramelot T A, Ovchinnikov S, Hao J, Bafna K, Norn C, Kang A, Bera A K, DiMaio F, Carter L, Chow C M, Montelione G T, and Baker D. De novo protein design by deep network hallucination. (2021) PMID: 34853475
2. Pesce F, Bremer A, Tesei G, Hopkins J B, Grace C R, Mittag T, and Lindorff-Larsen K. Design of intrinsically disordered protein variants with diverse structural properties. (2024) PMID: 39196930
3. Tretyachenko V, Vymětal J, Bednárová L, Kopecký V Jr, Hofbauerová K, Jindrová H, Hubálek M, Souček R, Konvalinka J, Vondrášek J, and Hlouchová K. Random protein sequences can form defined secondary structures and are well-tolerated in vivo. (2017) PMID: 29133927
4. Hunter K, Brandt T, Guadalupe K, Kolamunna K C, Lotthammer J M, Shamoon N M, Niblo J K, Nicholson B, Day L M, Martinez A, Holehouse A S, Sukenik S, and Emenecker R J. Rational design of disordered proteins for sequence-function investigation. (2026) PMID: 42527600
5. Lambert S A, Jolma A, Campitelli L F, Das P K, Yin Y, Albu M, Chen X, Taipale J, Hughes T R, and Weirauch M T. The human transcription factors. (2018) PMID: 29425488
